# Alteration of myocardial structure and function in RAF1-associated Noonan syndrome: Insights from cardiac disease modeling based on patient-derived iPSCs

**DOI:** 10.1101/2022.01.22.477319

**Authors:** Saeideh Nakhaei-Rad, Farhad Bazgir, Julia Dahlmann, Alexandra Viktoria Busley, Marcel Buchholzer, Fereshteh Haghighi, Anne Schänzer, Andreas Hahn, Sebastian Kötter, Denny Schanze, Ruchika Anand, Florian Funk, Andrea Borchardt, Annette Vera Kronenbitter, Jürgen Scheller, Roland P. Piekorz, Andreas S. Reichert, Marianne Volleth, Matthew J. Wolf, Ion Cristian Cirstea, Bruce D. Gelb, Marco Tartaglia, Joachim Schmitt, Martina Krüger, Ingo Kutschka, Lukas Cyganek, Martin Zenker, George Kensah, Mohammad R. Ahmadian

## Abstract

Noonan syndrome (NS), the most common among the RASopathies, is caused by germline variants in genes encoding components of the RAS-MAPK pathway. Distinct variants, including the recurrent Ser257Leu substitution in RAF1, are associated with severe hypertrophic cardiomyopathy (HCM). Here, we investigated the elusive mechanistic link between NS-associated RAF1S257L and HCM using three-dimensional cardiac bodies and bioartificial cardiac tissues generated from patient-derived induced pluripotent stem cells (iPSCs) harboring the pathogenic RAF1 c.770C>T missense change. We characterize the molecular, structural and functional consequences of aberrant RAF1 –associated signaling on the cardiac models. Ultrastructural assessment of the sarcomere revealed a shortening of the I-bands along the Z disc area in both iPSC-derived RAF1S257L cardiomyocytes, and myocardial tissue biopsies. The disease phenotype was partly reverted by using both MEK inhibition, and a gene-corrected isogenic RAF1L257S cell line. Collectively, our findings uncovered a direct link between a RASopathy gene variant and the abnormal sarcomere structure resulting in a cardiac dysfunction that remarkably recapitulates the human disease. These insights represent a basis to develop future targeted therapeutic approaches.

## Introduction

Germline variants of genes encoding RAS-mitogen-activated protein kinase (MAPK) signaling components result in a set of developmental disorders collectively referred to as RASopathies. They are characterized by craniofacial dysmorphism, delayed growth, variable neurocognitive impairment, and cardiac defects, including hypertrophic cardiomyopathy (HCM). HCM is known to vary in prevalence and severity depending on the genotype [3, 19, 52, 55]. At the organ level, HCM is manifested as an increase in the left ventricular wall thickness that may result in diastolic dysfunction, outflow obstruction, increased risk of heart failure, stroke and cardiac arrhythmia [38].

In Noonan syndrome (NS), the most prevalent disease entity among the RASopathies, development of HCM is associated with significant morbidity and risk of cardiac death. Notably, more than 90% of NS individuals with a *RAF1* variant are affected by HCM, whereas the overall frequency of HCM in NS is only about 20% [47, 53]. One variant, c.770C>T (p.Ser257Leu), accounts for more than 50% of cases with *RAF1*-associated NS (NSEuroNet database). Similar to other variants affecting transducers and positive modulatory proteins with role in the MAPK signaling cascade, the Ser257Leu amino acid substitution confers a functional enhancement (gain-of-function) to RAF1 kinase activity [38, 54]. Functional studies on RAF1 activity have mainly been focused on its role in activation of the MAPK pathway in cancer [8]. Recently, new paths of RAF1 cardiac-specific signaling towards ERK5 and calcineurin-NFAT have been discovered [9, 29, 59]. However, how coaction of RAF1-specific signaling pathways in cardiomyocytes (CMs) results in the phenotypical changes that lead to the development of HCM remains an open question. At the cellular level, HCM is characterized by an increase in CM size, re-activation of the fetal gene program, change in the amplitudes of calcium (Ca^2+^) transients, increased protein synthesis, and changes in the organization of the sarcomere structure and dysfunctional contractility [46]. Therefore, there is a critical need to combine signaling, phenotypical, and functional studies to dissect and understand the pathogenesis of HCM driven by RAF1^S257L^ gain-of-function (GoF).

To analyze the cardiac-specific function of RAF1 signaling, pure populations of human CMs are required to establish informative *in vitro* HCM disease models, where CMs should be able to respond to external stimuli and stress conditions as well as increase cell size rather than cell number. CMs generated from patient-specific induced pluripotent stem cells (iPSCs) have been used as human HCM models of several genetic etiologies as well as for drug screening [34, 40, 42, 66, 67]. Such *in vitro* models may recapitulate several HCM features at the cellular level, including the increase in cell size, aberrant calcium handling, reactivation of the fetal gene programs, and arrhythmia [13, 26, 29, 34, 59].

In this study, we generated and used two different three-dimensional (3D) human cardiac cell models, cardiac bodies (CBs) and bioartificial cardiac tissues (BCTs) to elucidate the molecular and cellular mechanism of RAF1-induced HCM. To this end, two iPSC lines derived from two different individuals carrying the recurrent *RAF1* c.770C>T (p.Ser257Leu) variant and three independent control iPSC lines were used (Fig. 1A). In iPSC-derived cardiomyocytes, we elucidated the effects of heterozygous RAF1^S257L^ on sarcomere structure, contractile behavior, Ca^2+^ handling, and intracellular signaling. Treatment with a MEK inhibitor (PD0325901) rescued the observed HCM phenotype caused by RAF1 GoF, which was validated using a gene-corrected isogenic iPSC line. Importantly, critical structural findings of the *in vitro* models matched the findings of a myocardial biopsy of one of the patients.

**Figure 1.**
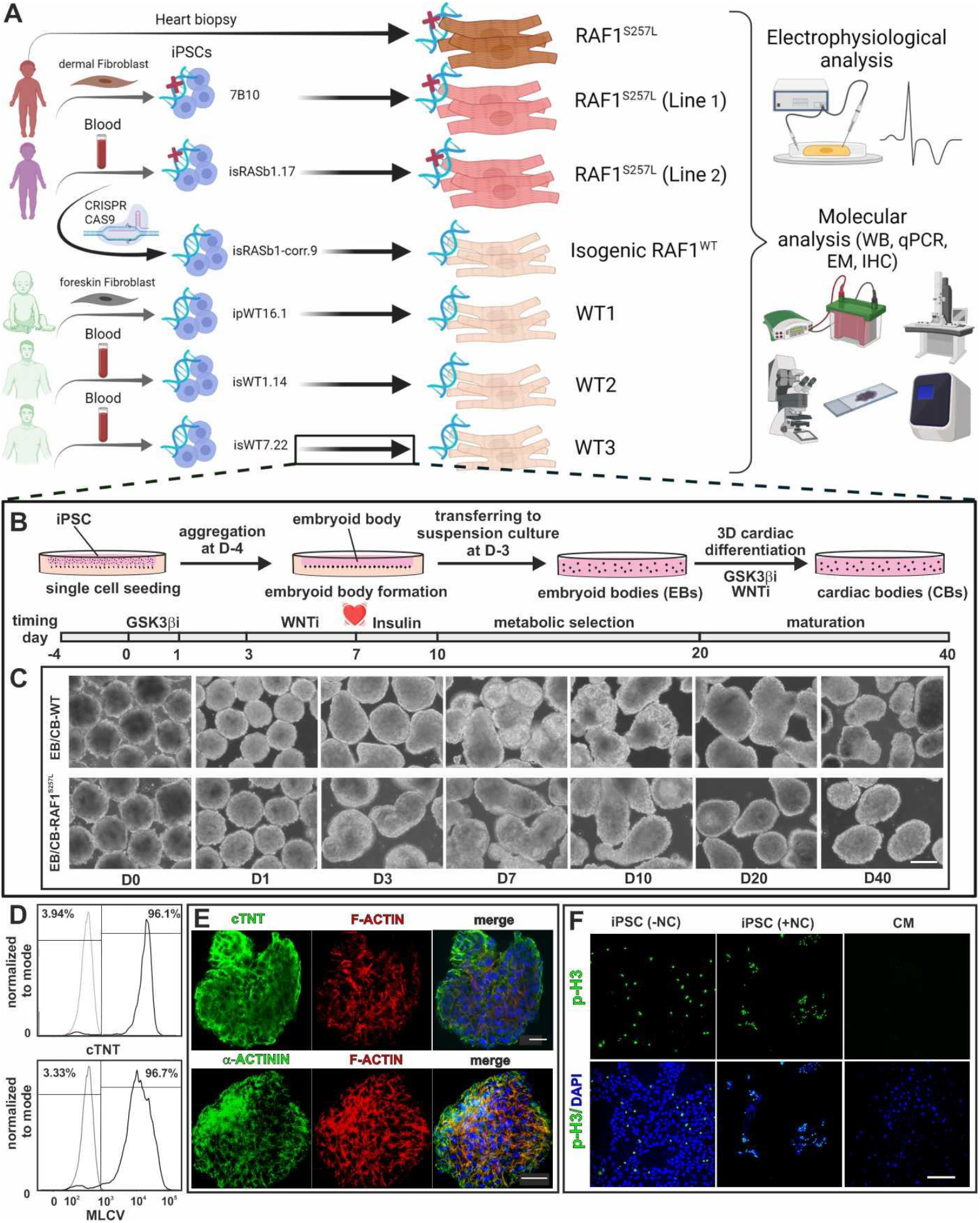
Study overview and details of 3D cardiac differentiation of iPSCs with WNT signaling modulation. A) Summary of the donor cells and the iPSC lines together with an overview of different analysis approaches used in this study. B) Schematic overview of embryoid body (EB) formation using agarose microwells combined with the stages and time lines of EBs differentiation to cardiac bodies (CBs). C) Light microscopic pictures of EBs/CBs during the course of cardiac differentiation and metabolic selection. Scale bar, 200 μm. D) Exemplary histograms of flow cytometric analysis of dissociated CBs displayed efficient cardiac differentiation towards ventricular cardiomyocytes by analysis of MLCV2 and cTNT positive cells (RAF1^S257L^-1). Isotype controls depicted in light gray. E) Immunofluorescence staining of a representative CB for cTNT and α-actinin expression (RAF1^S257L^-1). Scale bar, 20 μm. F) Illustration of mitotic cells stained with the mitotic marker p-H3 in iPSC cells and CMs. CMs showed no proliferative behavior as compared to iPSCs, which were arrested in mitosis by 100 nM nocodazole (NC) treatment. Scale bar, 200 μm.

## Results

### Generation of patient-specific iPSCs

To model the RAF1-induced HCM based on patient-derived iPSCs, two NS individuals with severe HCM, heterozygous for the pathogenic *RAF1 c.770C>T* variant (p.Ser257Leu; RAF1^S257L^ hereafter), were recruited for this study (Fig. 1A). From both individuals, patient-specific iPSC lines were generated by integration-free methods (Fig. S1 and S2). Reprogramming of fibroblasts to iPSC was performed using a cocktail of transcription factors *OCT3/4, SOX2, L-MYC, TRP53*, and *EBNA-1* as previously described [44]. Furthermore, an isogenic iPSC line from one of these individuals was generated via CRISPR/Cas9 technology (Fig. S2H). In parallel, control iPSC lines from three healthy individuals (iPSC-WT1,2,3) were used and characterized (Fig. S3)[58]. The iPSC-WT and iPSC-RAF1^S257L^ lines were positive for different pluripotency markers, including alkaline phosphatase (Fig. S1B and S3B), OCT4, TRA-1-60, and SSEA4 (Fig. S1C, S2A-C and S3C). Sequencing verified the presence of the heterozygous *RAF1* c.770C>T variant in the iPSC-RAF1^S257L^ lines (Fig. S1D and S2D), and karyotyping confirmed the stable chromosomal integrity of the iPSC lines (Fig. S1E and S2E). Spontaneous embryoid body (EB) formation and spontaneous undirected differentiation confirmed the tri-lineage differentiation potential of each line (Fig. S1F-G, S2F-G and S3F-G).

### Cardiac differentiation of patient-specific iPSCs

Human iPSC lines were differentiated into cardiomyocytes in 3D aggregates by temporal modulation of canonical WNT signaling as described previously [5, 35]. After EB formation (Fig. 1B), differentiation was initiated with GSK3β-inhibitor CHIR-99021 for 24h followed by WNT pathway inhibitor IWR-1 for 48h (Fig. 1C). Five to seven days after induction of cardiac differentiation, CBs started to contract spontaneously. At d10 of differentiation, metabolic selection was performed by replacing glucose with lactate in the medium, leading to a purified population of CBs taking advantage of the unique metabolic pathways and energy sources characterizing CM (Fig. 1C). Analyses showed that the cells were negative for the presence of pluripotency markers at mRNA and protein levels (data are not shown) and positive for cardiac markers (Figs. 1D,E and S5A). At d40 of differentiation, CBs were more than 96% cTNT-positive and 96% MLC2v-positive, as analyzed by flow cytometry (Fig. 1D). Immunofluorescence of cryosections of CBs revealed homogenous populations of cTNT- and α-actinin-positive cells in the center as well as in the periphery of the CBs (Fig. 1E).

Cardiac hypertrophy is defined as enlargement of the cardiomyocytes. iPSC-derived CMs were tested for their inability to proliferate in response to external stimuli. To this goal, d40 CBs were dissociated, and single CMs were seeded on coverslips. Thereafter, CMs were fixed and assessed for the presence of the mitotic marker phospho-Ser10-histone 3 (p-H3), confirming their negative p-H3 staining at d47 (Fig. 1F). As a positive control, proliferative human iPSCs were treated for 12 h with 100 nM nocodazole (NC) to be arrested in mitosis (Fig. 1F and S4A,B). Cell cycle analysis indicated that NC-treated human iPSCs were mainly captured at the G2/M phase and stained positive for p-H3 (Fig. S4B,C). Moreover, dissociated CMs at d40 were treated with 100 μM L-phenylephrine (PE), an α-adrenergic agonist, for 7 days. These cells remained p-H3 negative, but increased in size (Fig. S4D-F), which is consistent with PE’s pro-hypertrophic activity. Collectively, these data indicate that in our experimental system employed iPSC-derived CMs had the ability to increase in size in response to hypertrophic stimuli, but did not undergo proliferation (hyperplasia), providing a suitable *in vitro* model to study the molecular mechanism implicated in RAF1-driven HCM.

### Signaling events in RAF1^S257L^ CBs

Serine^257^ (S257) is located in close proximity to the negative regulatory phosphorylation site S259 within the conserved region 2 (CR2) of RAF1 (Fig. 2A). The latter provides a docking site for 14-3-3 proteins, thereby stabilizing RAF1 in its auto-inhibited state [14, 54]. To determine the impact of the S257L substitution on the S259 phosphorylation status in cells with endogenous expression of the kinase, we immunoprecipitated total RAF1 protein with an anti-RAF1 antibody from lysates of undifferentiated iPSCs-WT and iPSCs-RAF1^S257L^(line1). Immunoblot analysis with anti-RAF1 and anti-p-RAF1^S259^ revealed up to 44% reduction in the levels of RAF1^S257L^ phosphorylated protein, compared to WT RAF1 at S259 (Fig. 2B). Therefore, due to the heterozygous status of the mutation in the model system, it can be assumed that the majority of the mutant RAF1 protein remains unphosphorylated, accounting for almost 50% of total RAF1. This observation demonstrates a reduced ability of RAF1^S257L^ to be subjected to the 14-3-3 inhibitory control at physiological conditions.

**Figure 2.**
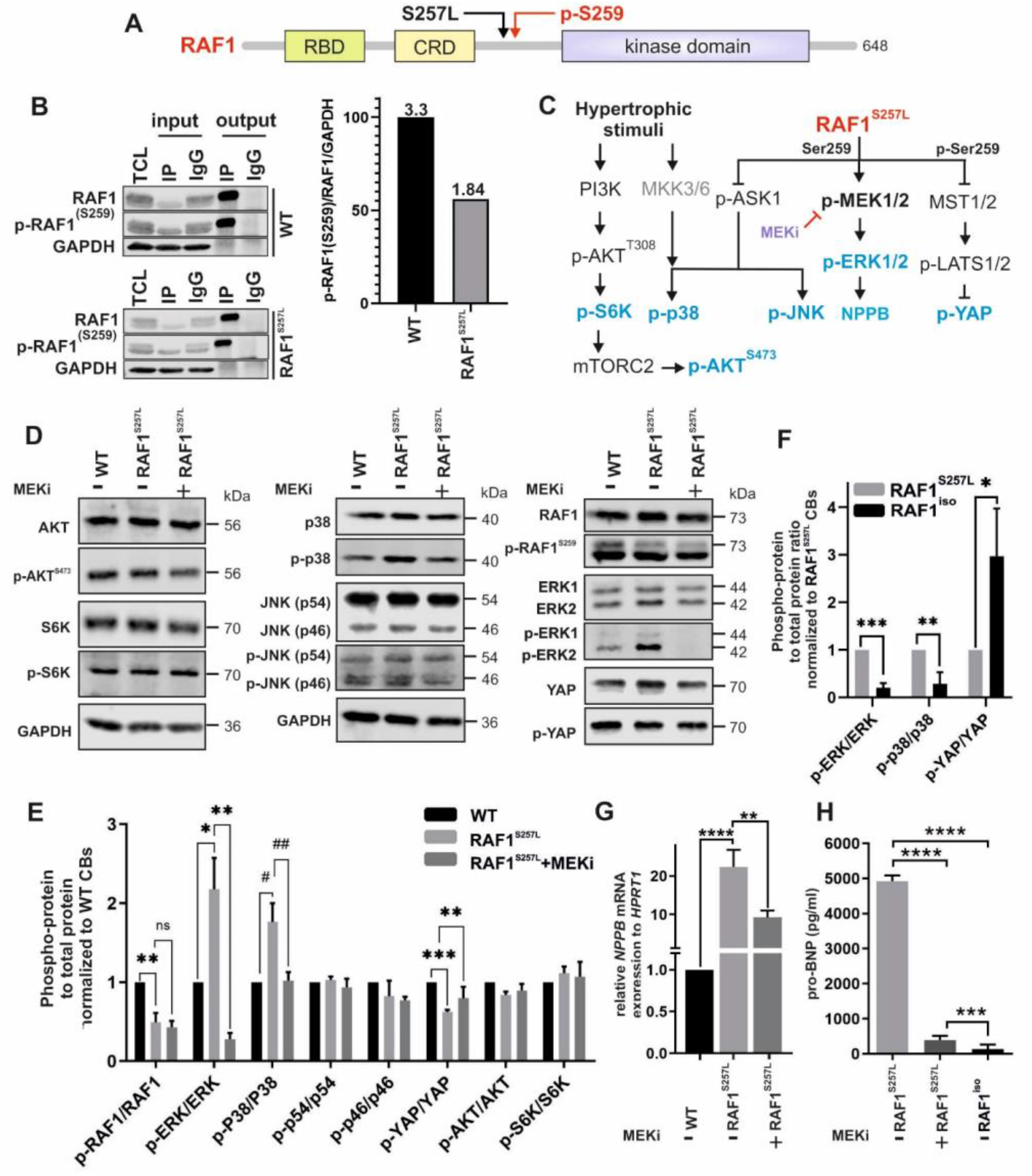
The effect of the RAF1^S257L^ variant on the activity of selected effector kinases downstream of RAF1. A) Domain organization of RAF1 kinase with the typical functional domains, including the RAS-binding domain (RBD), the cysteine-rich domain (CRD), and the kinase domain. The adjacent sites of S257L variant and the inhibitory S259 phosphorylation (p-S259) are highlighted. B) Immunoprecipitation and quantification of total and p-RAF1^S259^ in WT and RAF1^S257L^ iPSCs. Total RAF1 was pulled down from lysates of WT and RAF1^S257L^ iPSCs using an anti-RAF1 specific antibody. IgG was applied as an isotype control. Immunoblotting was carried out using anti-RAF1 and anti-p-RAF1^S259^ antibodies. For quantification, signal intensities of p-RAF1^S259^ were divided by those for total RAF1. GAPDH was used as a loading control. TCL, total cell lysate; IP, immunoprecipitation; IgG, Immunoglobulin G. C) Schematic diagram summarizing the signaling molecules investigated downstream of hypertrophic stimuli and RAF1. Proteins marked in blue letters were investigated at the protein level by immunoblotting. D) Representative immunoblots of p-AKT *vs*. AKT, p-S6K *vs*. S6K, p-RAF1^259^ *vs*. RAF1, p-ERK1/2 *vs*. ERK1/2, p-YAP *vs*. YAP, p-p38 *vs*. p38, and p-JNK *vs*. JNK using cell lysates from WT and RAF1^S257^ CBs (d24). E) Phospho-protein *vs*. total protein ratio quantification as shown in D. *P < 0.05, **P < 0.01, ***P < 0.001, ****P < 0.0001, unpaired 2-tail t-test. # P < 0.05, ## P < 0.01, unpaired 1-tail t-test. n≤2. F) Phospho-protein *vs*. total protein ratio quantification of western blot results for selected pathways in RAF1^S257L^ CBs vs. gene corrected line, RAF1^iso^ CBs (d24). *P < 0.05, **P < 0.01, ***P < 0.001, unpaired 2-tail t-test. n≤2. G) qPCR analysis of *NPPB* transcription levels. **P < 0.01, ****P < 0.0001, unpaired 2-tail t-test, n=3. H) ELISA analysis of pro-BNP levels released in the cell culture supernatant of CB’s (pg/ml). ***P < 0.001, ****P < 0.0001, unpaired 2-tail t-test. n=8.

Next, we investigated the activity of selected RAS/RAF-dependent signaling in WT and RAF1^S257L^ CBs in the presence and absence of the MEK inhibitor PD0325901 (MEKi; Fig 2C). In untreated CBs, the PI3K-AKT-S6K-mTORC-AKT and RAF1-ASK1-JNK signaling axes did not show remarkable differences between RAF1^S257L^ and control CBs (Fig. 2D,E). However, increased levels of p-ERK1/2 and p-p38, and decreased levels of p-YAP in RAF1^S257L^ CBs were documented. The p-ERK1/2 and p-p38 levels were significantly reduced upon treatment with MEKi, while a significant increase in the level of p-YAP was observed (Fig. 2D,E and S5B,C). Furthermore, we examined the impact of the signaling signature of CRISPR-corrected RAF1^iso^ CBs (RAF1^L257S^) *vs*. its mother clone (RAF1^S257L^ CBs) and found the opposing pattern of phosphorylation of the former pathways (Fig 2F), which highlights the explicit impact of the RAF1 point mutation at Serine257 on the observed signaling patterns.

Quantitative real-time PCR (qPCR) analysis showed significant upregulation of *NPPB* in RAF1^S257L^ CBs, which was partially reverted in the presence of MEKi (Fig. 2G and S5D). The gene *NPPB* encodes for BNP (brain natriuretic peptide), a well-known clinical biomarker for heart failure and is upregulated during hypertrophy due to a return to a fetal-like gene expression program. Next, we examined the levels of secreted pro-BNP in medium of the cultured CB variants. Interestingly, the results indicated that RAF1^S257L^ CBs secrete more than 12 and 30-fold amounts of BNP in their medium compared to MEKi-treated and isogenic control CBs, respectively (Fig 2H). Notably, the isogenic control elucidated even lower amount of BNP than MEKi treated cells, which may indicate for activity of the parallel pathways beside RAF1-MAPK in regulation of BNP levels, *e.g*., p-RAF1-MST2-YAP (Fig. 7).

### RAF1^S257L^ alters stretch-shortening of sarcomere

Familial non-syndromic HCM caused by mutations in sarcomeric proteins is known to affect sarcomere architecture [45]. To address the question of whether the sarcomeric architecture is also altered in RAF1^S257L^-associated HCM, CBs at d40 were dissociated and single CMs were cultured for 7 d on coverslips. RAF1^S257L^ CMs showed less oriented and more disarrayed myofilaments as compared to WT CMs (Fig. 3A), which was confirmed by immunohistochemistry (IHC) and electron microscopy (EM). In particular, RAF1^S257L^ CBs revealed shortened or more contracted I-band regions and thickened Z-line pattern (Fig. 3B). Left ventricular cardiac tissue (CT) was available for one of the NS individuals with RAF1^S257L^, and staining of desmin and cardiac troponin I confirmed a disorganized sarcomeric structure with shortened I-bands, as seen on iPSC-derived CMs and CBs along with a thickened Z-line (Figs. 3C,D). Notably, RAF1 was localized and condensed near the sarcomeric structures in RAF1^S257L^ CMs, while in WT CMs RAF1 expression was more cytoplasmic (Fig. 3E).

**Figure 3.**
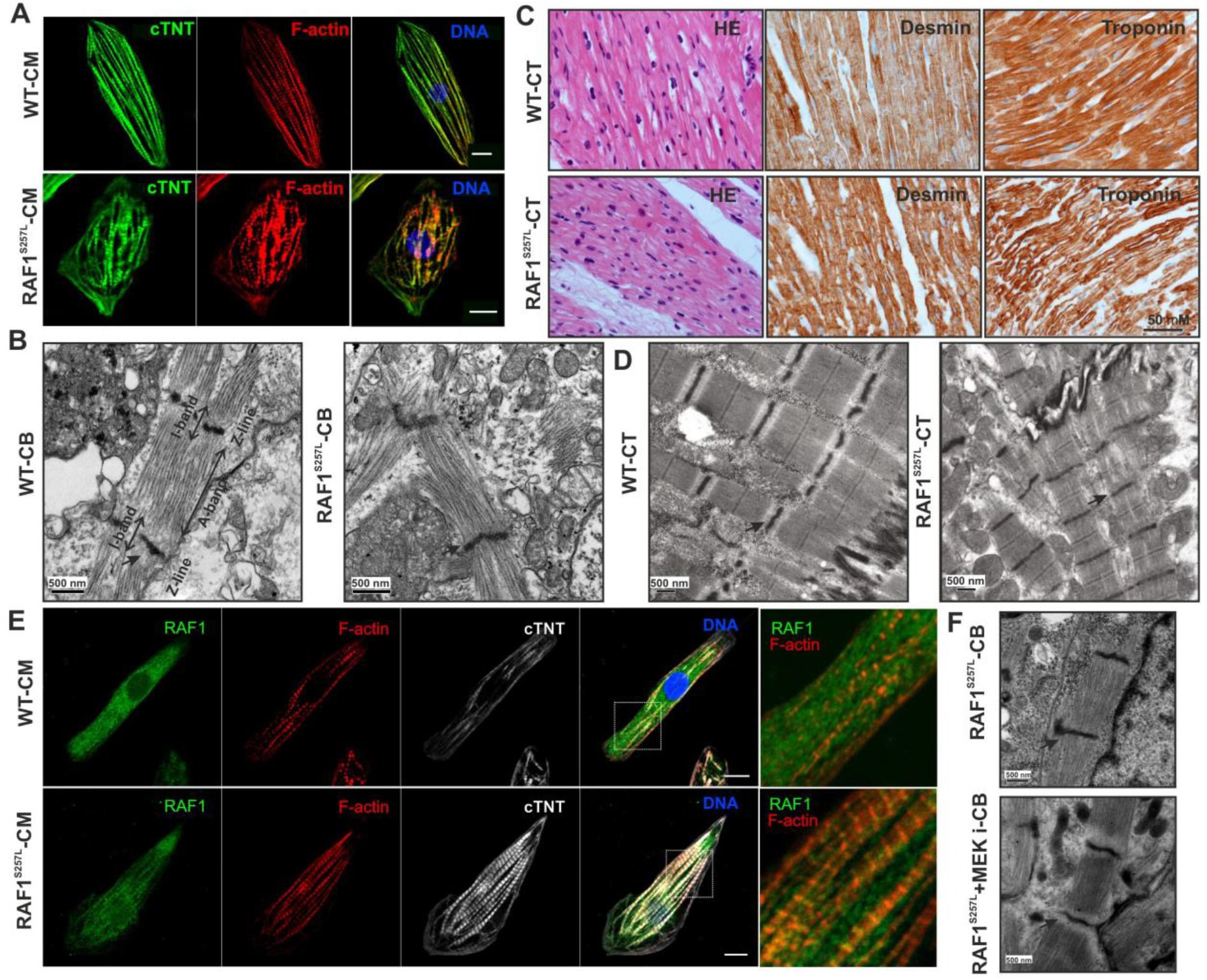
Aberrant RAF1^S257L^ activity impairs the cytoarchitecture of human iPSC-derived cardiomyocytes. A) Dissociated cardiac bodies were seeded on Geltrex-coated coverslips for 7 d and stained for cTNT and F-actin (Scale bar, 10 μm). B) Representative EM images from RAF1^S257L^ CBs revealed a stronger myofibrillar disarray accompanied by shortened I-bands and a thickened Z-line pattern as compared to WT CBs. C) IHC analysis of RAF1^S257L^ CTs from one of the NS individuals with *RAF1 c. 770C>T* variant for desmin and troponin showed myofilament disarray. D) Representative EM images of the same RAF1^S257L^ CTs as in C exhibited shortened I-bands and a thickened Z-line pattern consistent with RAF1^S257L^ CBs in B. E) Representative ICC images of RAF1^S257L^ and WT CMs at d90 post-differentiation showed RAF1 co-localization with cTNT and F-actin at the sarcomere (Scale bar, 10 μm). F) EM images of RAF1 mutated RAF1^S257L^ CBs (d40) treated with 0.2 μM MEKi from d12 of differentiation.

To examine the influence of the dysregulated MAPK signaling on the observed phenotype, we treated the RAF1^S257L^ CBs with PD0325901 (0.2 μM) at early stages of development (d12) until d40. Remarkably, ultrastructure analysis revealed that MEKi treatment restored the I-band width around the Z-line in the RAF1^S257L^ CBs (Fig. 3F and S5H). To confirm the observed I-band shortening, we prepared cryosections of RAF1^S257L^ and WT BCTs and dissected a part of giant sarcomeric protein titin by immunostaining the PEVK domain of titin to mark the I-band region and α-actinin staining to indicate Z-lines (Fig. 4A). PEVK domains of titin and α-actinin were strikingly co-localized in RAF1^S257L^ BCTs, while clearly separated from each other in WT BCTs (Fig. 4A and B), suggesting a major shortening of the I-band region with dislocation of the titin PEVK region in the RAF1^S257L^ BCTs. Area histograms of the selected sub-images (Fig. 4B; boxed and magnified) were created to determine the distance between the two PEVK segments and quantify the average distances of more than 50 different Z-lines per sample. These abnormalities of the overlapped peaks corresponding to the two PEVK domains relative to α-actinin were remarkably restored and comparable to WT BCTs when RAF1^S257L^ BCTs were treated with MEKi (Fig. 4C). The significant separation of the distance between the two adjacent PEVK segments upon MEKi treatment strongly suggests a functional normalization of the sarcomeric structures in RAF1^S257L^ BCTs (Fig. 4D). Moreover, by measuring the average distance between α-actinin signals as the Z-line marker, a decrease in the sarcomere lengths was observed for the RAF1^S257L^ BCTs compared to WT BCTs (Fig. 4E). We further determined a significant increase in the mRNA levels of the predominant longer and more compliant (*N2BA*) titin isoform towards the shorter and stiffer isoform (*N2B*) in RAF1^S257L^ BCTs (Fig. 4F).

**Figure 4.**
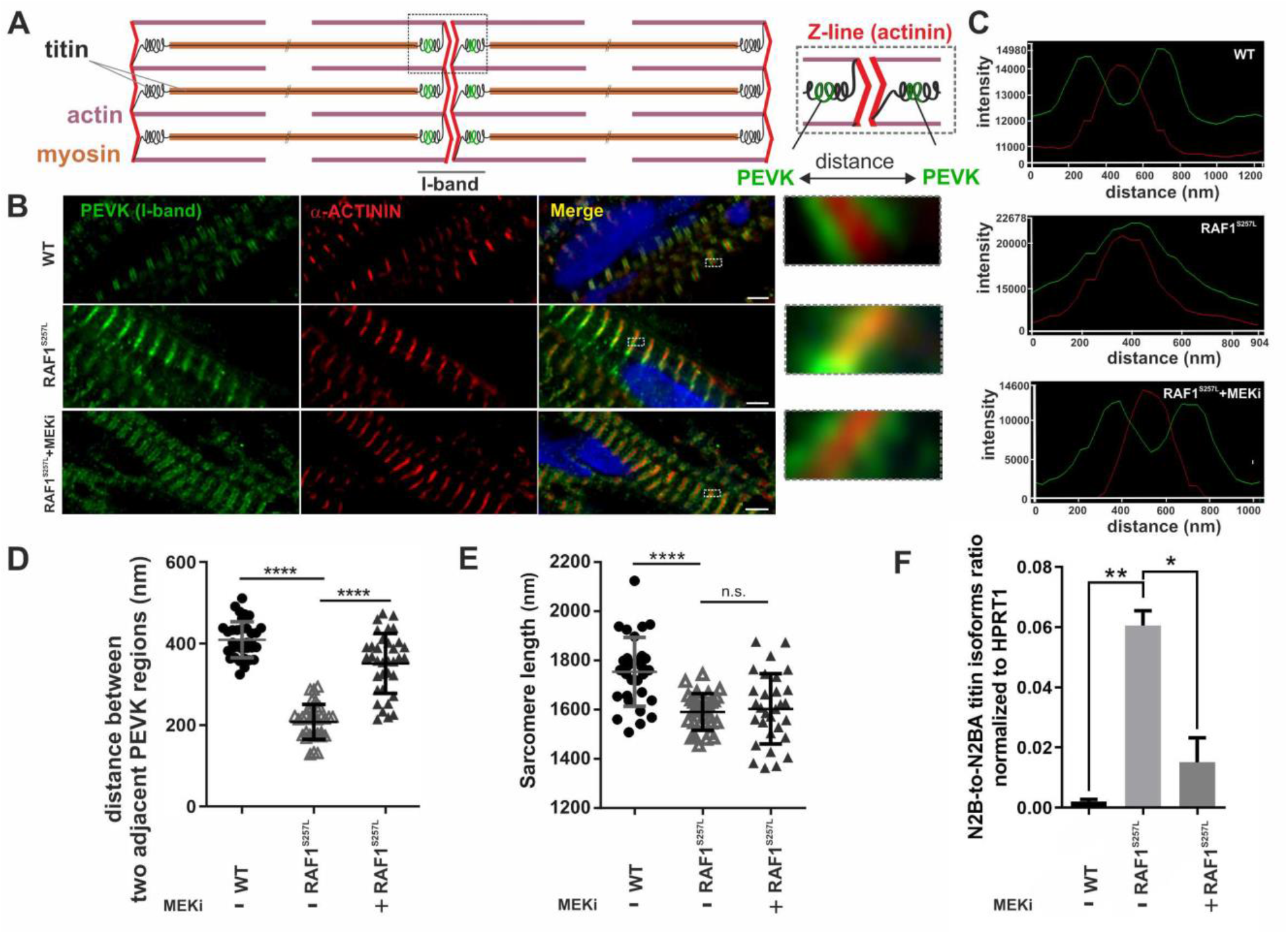
Hyperactive RAF1^S257L^ triggers a shorter I-band phenotype A) Schematic view of the sarcomeric organization. B) IHC analysis of 8-μm cryosections of WT and RAF1^S257L^ BCTs with PEVK segment of titin’s I-band, α-actinin 2 as the Z-line marker, and DAPI for DNA staining. C) Histogram of selected boxes on G were imported base on the intensity and overlaps of emitted fluorescent lights using the Zeiss LSM 880 Airyscan confocal microscope software. D) Averaged distance (nM) between two adjacent PEVK segments was measured for more than 50 different sarcomeric units for each condition and statistically evaluated. ****P < 0.0001, unpaired 2-tailed t-test. E) Averaged sarcomere length (nM) was measured for more than 50 different sarcomere units for each condition by measuring the distance between two parallel Z-lines (α-actinin) and statistically evaluated. ****P < 0.0001, unpaired 2-tailed t-test. F) qPCR analysis of ratio of the *N2B*-to-*N2BA* titin isoforms expression levels in CBs. *P < 0.05, **P < 0.01, unpaired 2-tailed t-test.

Collectively, the data demonstrate that RAF1 GoF promotes I-band shortening and reduces the flexibility of the spring elements of the titin I-band region.

### Aberrant expression of sarcomeric regulatory proteins

The impaired sarcomere organization of RAF1^S257L^ CMs prompted us to quantitatively analyze the expression of key sarcomeric components, including troponins, myosins, and actin-related proteins. In comparison to WT CBs at d24, RAF1^S257L^ CBs strikingly exhibited higher levels of *MYH7* and *MYL2*, but lower levels of *TTN, MYH6, MYL7*, and *α-SMA* (Fig. 5A). We analyzed the *MYH7*-to-*MYH6* ratio in CBs at two different maturation stages (d24 and d47). Both, immature (d24) and more mature (d47) CBs displayed a significant increase in the *MYH7*-to-*MYH6* ratio (Fig. 5B and S5E). Notably, MEKi treatment partially reversed the *MYH7*-to-*MYH6* ratio in d47, but not d24, RAF1^S257L^ CBs.

**Figure 5.**
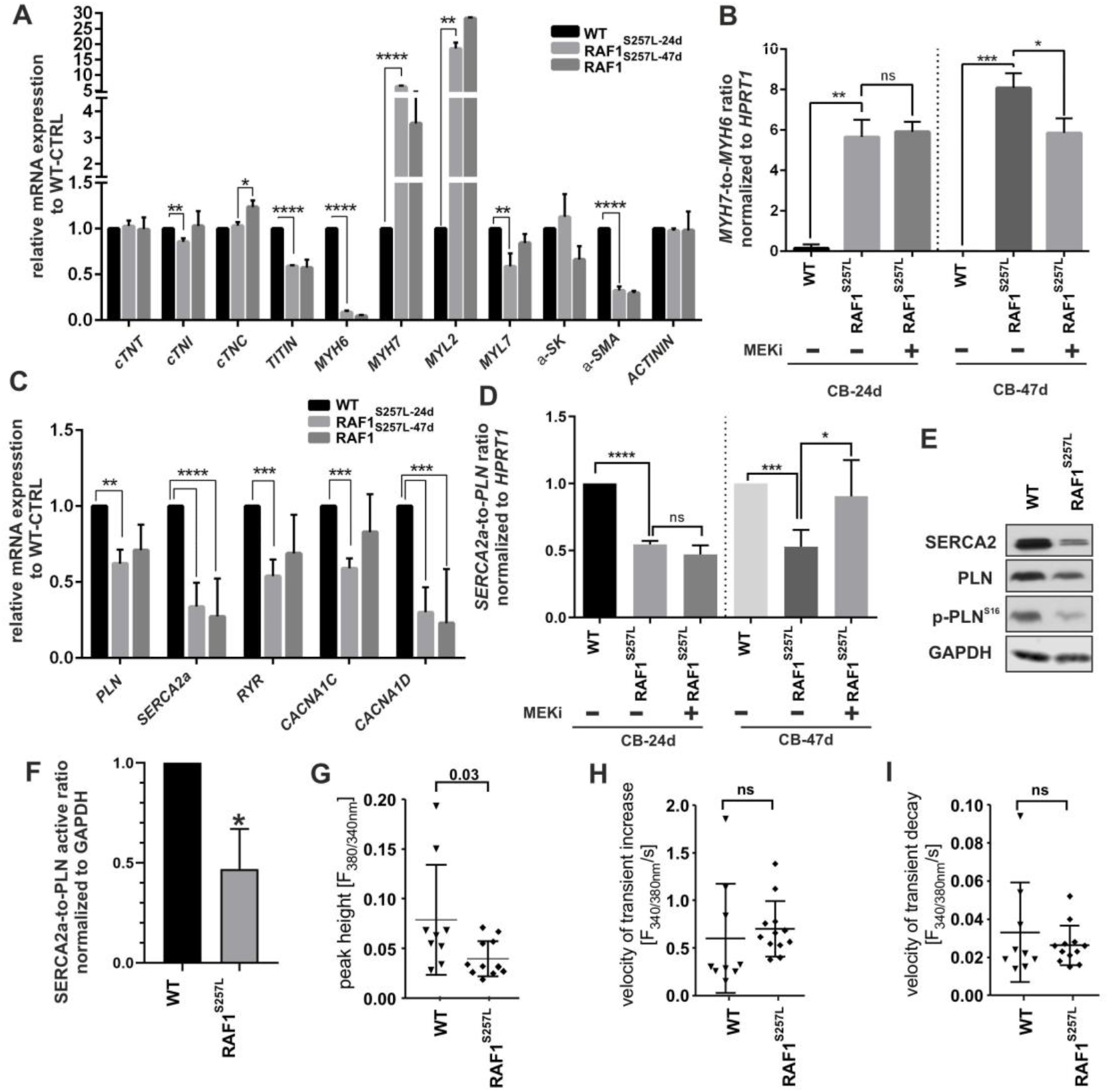
Abnormal expression of proteins involved in sarcomere function and calcium handling. The experiments in A-F were conducted with WT CBs, RAF1^S257L^ CBs, and RAF1^S257L^ CBs treated with 0.2 μM MEKi from d12 of differentiation. The data are averaged from three independent experiments in biological triplicates. *P < 0.05, **P < 0.01, ***P < 0.001, ****P < 0.0001, unpaired 2-tail t-test. A) qPCR analysis of mRNAs related to sarcomere proteins. B) *MYH7*-to-*MYH6* ratio. C) qPCR analysis of mRNAs related to regulation of calcium transients. D) *SERCA2a*-to-*PLN* ratio. E) Immunoblot analysis of SERCA2, PLN, and p-PLN^Ser16^ in CBs at d24. F) The ratio of SERCA2 to PLN was calculated by measuring the ration of SERCA2 to PLN/p-PLN^Ser16^ G-I) Ca^2+^ transients were measured in Fura2-loaded CMs and expressed as the ratio of fluorescence emission at 340 nm and 380 nm. Bar graphs display the peak height of Ca^2+^ transients (G) and the velocities of cytosolic Ca^2+^ increase (H) and decrease (I). Each data point represents the average of 10 transients obtained from a single CM. Nine wild-type and twelve RAF1^S257L^ CMs were analyzed in total.

### Reduced Ca^2+^ transients RAF1^S257L^ CMs

Next, we investigated a possible dysfunctional calcium handling of RAF1^S257L^ CBs, which is a central feature of HCM [34]. We first analyzed the expression of the components that regulate intracellular calcium cycling in WT and RAF1^S257L^ CBs at d24 and d47. These components include ryanodine receptor type-2 (RyR2), sarco/endoplasmic reticulum Ca^2+^-ATPase (SERCA2A), phospholamban (PLN), and the calcium voltage-gated channel alpha (CACNA) subunits 1C and 1D (L-type calcium channels (LTCCs)). At d24, *PLN*, *SERCA2A*, *RYR*, and *CACNA1C* mRNA expression was downregulated in RAF1^S257L^ CBs (Fig. 5C). Furthermore, *SERCA2A* and *CACNA1D* expression was significantly reduced in RAF1^S257L^ CBs at d47 (Fig. 5C). Remarkably, the decrease of the *SERCA2-to-PLN* ratio in RAF1^S257L^ CBs was reversed to WT levels upon MEKi treatment in d47 CBs (Fig. 5D and S5F).

Phosphorylation of PLN at Ser^16^ inhibits PLN activity. SERCA2a, PLN, and p-PLN^Ser16^ were reduced in RAF1^S257L^ compared to WT, CBs (Fig 5E). The SERCA2/active PLN ratio, which was calculated at protein levels by measuring the ratio of SERCA2 to PLN/p-PLN^Ser16^, was also significantly reduced in RAF1^S257L^ CBs (Fig. 5F). The changes in SERCA2a and PLN expression and the ratio of PLN to p-PLN^Ser16^ were consistent with aberrant calcium handling properties, a characteristic of maladaptive hypertrophy [11, 34]. Therefore, we measured intracellular calcium transients by seeding CMs (d47) on Geltrex-coated coverslips and loaded the cells with the calcium indicator Fura-2. The calcium release was significantly decreased in RAF1^S257L^ CMs as compared to WT CMs (Fig 5G). Despite reduced transients, the kinetics of cytosolic calcium rise and decrease in RAF1^S257L^ CBs were not different from WT CBs (Fig. 5H and I). Collectively, these data suggest that the RAF1^S257L^ variant modulates the contractile cardiac function by impairing cellular calcium cycling.

### Negative impact of RAF1^S257L^ on the contractile apparatus

Next, we analyzed the effect of RAF1^S257L^ on cellular contractility using a multimodal bioreactor system to generate, cultivate, stimulate, and characterize BCTs non-invasively [32]. RAF1^S257L^ BCTs showed a significantly higher spontaneous contraction frequency as compared to WT BCTs, which was significantly reduced by 0.1 and 0.2 μM MEKi (Fig 6A). Quantification of cross-sectional areas, which is a measure of myocardial thickening, showed no significant difference between WT and RAF1^S257L^ BCT samples (Fig 6B). However, RAF1^S257L^ BCTs treated with MEKi had a reduced cross-sectional area. Active force and tension measurements revealed a significantly lower contractile force and contractile tensions for RAF1^S257L^ BCTs compared to WT BCTs (Fig 6C,D). RAF1^S257L^ BCTs treated with 0.1 μM MEKi had improved contractile force and re-established contractile tensions (Fig 6C,D). Moreover, analysis of morphology of contraction peaks revealed a longer time to peak and shorter time to 80% relaxation of RAF1^S257L^ BCTs compared to WT BCTs (Fig. 6E,F). Again, the latter was increased upon MEKi treatment. Notably, BCTs showed a much better rescue response to 0.1 μM MEKi treatment as compared 0.2 μM MEKi. In summary, RAF1^S257L^ BCT have higher contraction frequencies, less contractile force and tension, impaired contractile kinetics, and accelerated relaxation kinetics. Most of the observed effects were reversed by application of the MEKi, in a dose dependent manner.

**Figure 6.**
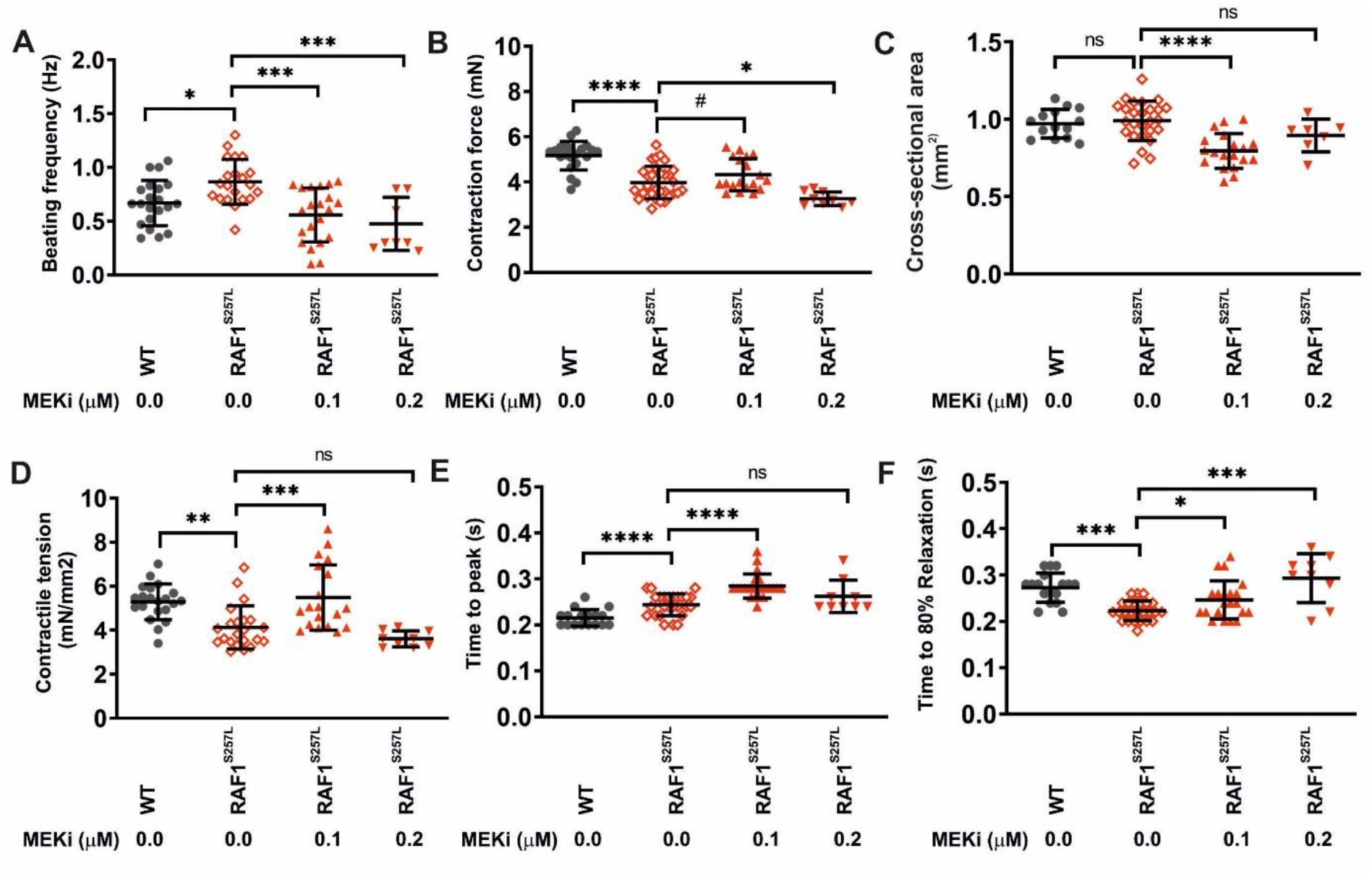
Aberrant contractility of RAF1^S257L^ BCTs and partial rescue by MEKi. Physiological measurements were conducted on day 24-28 of culture using WT, RAF1^S257L^ and RAF1^S257L^ +MEKi BCTs. N = 9-26 individual tissue samples per group. Depending on the presence of normally distributed values, one-way ANOVA or Kruskal-Wallis test was applied. *P<0.05, **P<0.01, ***P<0.001, ****P<0.0001. A) Spontaneous beating frequencies. B) Quantification of cross-sectional areas. C) Maximum contraction forces. D) Maximum contractile tensions based on the cross-sectional areas. E) Time to peak. F) Time to 80% relaxation.

**Figure 7.**
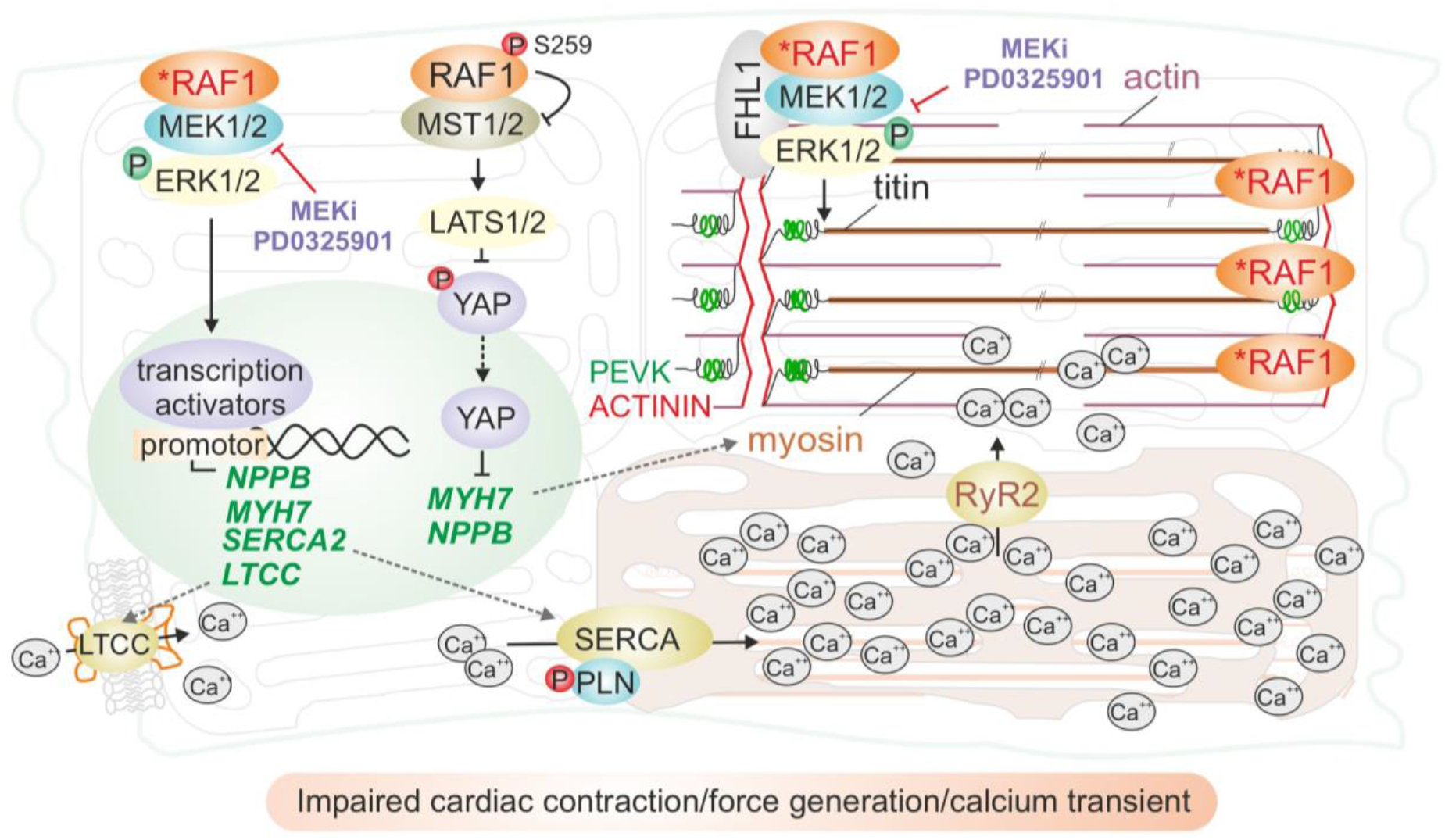
A proposed model of both *RAF1*-dependent cardiac signaling pathways, coupling calcium transients and contraction, and RAF1^S257L^-enhanced impairment of cardiac contraction, force generation, and calcium transients (for details, see “Discussion”).

## Discussion

The *in vitro* cellular reprogramming, differentiation, and tissue engineering of patient-derived samples reproducibly generated CBs and BCTs as human 3D disease models to investigate the molecular events contributing to HCM in RAF1-related Noonan syndrome. This genetically determined human disorder is caused by aberrantly enhanced function of the RAF1 kinase. We characterized the CBs’ and BCTs’ cytoskeletal and sarcomeric ultra-structures by super-resolution and EM imaging, to assess their calcium handling and contractility, and intracellular signal transduction. These complementary approaches identified reduced *MYH6* abundance over *MYH7*, elevated expression of *NPPB* and secretion of pro-BNP, decreased *SERCA2/PLN* ratio, sarcomeric fibril disarray, reduced force generation accompanied by a reduced rate of intracellular calcium transients, increased levels of p-ERK1/2, p-p38, and attenuation of p-YAP, as signatures of RAF1^S257L^ CMs. Most remarkably, RAF1^S257L^ CBs and BCTs as well as heart biopsy samples from the RAF1^S257L^ individuals revealed common ultrastructural features, namely shortened I-bands. The alterations in titin and shortened I-bands of RAF1^S257L^ CBs/BCTs were attenuated by treatment with the MEK inhibitor PD0325901. Collectively, our results suggest RAF1^S257L^-mediated activation of the MAPK pathway produces an abnormal cardiac phenotype involving structural and physiological aspects, that can be in part rescued with MEKi.

### RAF1^S257L^ signaling in human cardiomyocytes

CMs with heterozygous RAF1^S257L^ exhibited an approximately 50% reduced inhibitory phosphorylation of RAF1 at S259, which is consistent with previous reports [29, 47, 53]. The best studied RAF1 function is activation of the MEK1/2-ERK1/2 pathway, which regulates the activity of a wide range of signaling molecules in the cytoplasm and nucleus. Accordingly, higher p-ERK1/2 levels were detected in RAF1^S257L^ CMs compared to control CMs, also consistent with previous studies [9, 29] and, p-ERK1/2 levels were remarkably reduced in CBs upon MEKi treatment (Fig. 2D,E and S5B-C). In addition to its crucial role in normal heart development, the MAPK pathway may also act as central signaling node for many factors stimulating adaptive and maladaptive hypertrophy [18, 57]. However, the detailed mechanisms involving aberrant RAF1^S257L^-MEK1/2-ERK1/2 signal transduction to induce hypertrophy remain unclear. This phenomenon may be rooted in the regulation of cardiac specific-transcription activators or/and direct/indirect transcriptional modulations of cardiac components (Fig. 4F) via ERK1/2 regulation (nuclear substrates) as well as direct modulation of cardiac function by affecting contractile machinery (cytoplasmic/sarcomeric substrates; Fig. 7). Accordingly, we detected a differential expression of a fetal-like gene program, proteins of the contractile machinery, *MYH7* and, calcium transient regulators, in RAF1^S257L^ CBs as compared to WT and MEKi-treated CBs. The observed changes in the expression for these genes likely results from transcriptional activation of the hypertrophic-responsive gene promoters by GATA4, AP1, MEF2, NFAT, and NFκB, as described by previous studies [10, 25, 27]. Future investigations employing chromatin immunoprecipitation of these transcription factors from isolated nuclei of RAF1^S257L^ CBs combined with proteomic approaches may clarify the transcriptional regulation by MAPK pathway.

Activated RAF1 binds to MST2 (also called STK3) and inhibits the MST1/2-LATS1/2 pathway (Romano et al., 2010; Romano et al., 2014). As a consequence, YAP translocates into the nucleus, associates with TEAD to serve as a transcriptional co-activator, and regulates transcription of the mitogenic factors, including CTGF, NOTCH2 and c-MYC as well as miR-206. The MST1/2-LATS1/2-YAP axis is critical for heart development, growth, regeneration, and physiology [69]. It regulates proliferation in neonatal heart and growth and survival in adult heart [7, 65, 73]. The YAP-miR206-FOXP1 axis regulates hypertrophy and survival of CMs through upregulation of fetal genes, e.g., *MYH7* and *NPPB* [68]. We demonstrate here that the reduced inactivating phosphorylation of YAP in RAF1^S257L^ was restored upon MEKi treatment (Fig. 2D-F), which is likely explained by the enhanced RAF1^S257L^ kinase activity along with a switch of RAF1 binding from MST1/2 to MEK1/2 (Fig. 2C) [56]. We also observed increased expression of YAP targets, such as *MYH7* and *NPPB* in RAF1^S257L^ CBs. Highly elevated levels of the NPPB gene product pro-BNP were detected in the supernatant of RAF1^S257L^ CBs, considerably above the critical clinical thresholds defined for likelihood of a heart failure state in patients (Fig. 2H) [63]. This validates the observed signaling impact of hyperactive RAF1^S257L^ signaling on the fetal gene expression programs, directing the cells towards a heart failure condition. Considerably, we inhibited the MEK-MAPK axis in cardiac cells without targeting RAF1 directly, therefore we expected that only the MAPK-dependent phenotype is rescued upon MEKi treatment. However, we observed that parallel pathways downstream of RAF1 is also reverted by MEKi treatments such as MST2/YAP.

We propose that MEK inhibition abrogates the negative feedback phosphorylation of RAF1 by ERK, and thus restores RAF1 membrane localization and activity [12].

Other MAPKs besides ERK1/2, such as p38, JNK, and ERK5, appear to be involved in cardiac development, function, and also progression of myocardial disease (Jaffre et al., 2019a; Rose et al., 2010). Our data indicated an increase in p38 phosphorylation, but not JNK, in RAF1^S257L^ CBs. Aberrant p38 phosphorylation was reverted to (near) normal upon MEKi treatment. The molecular mechanism that underlies p38 activation *via* RAF1^S257L^ in cardiac cells is unclear. We propose that elevated levels of p-p38 in RAF1^S257L^ cells may be a compensatory response of cells to reduce the effects of the sustained ERK activity towards an unknown ERK-dependent or ERK-independent positive feedback regulation of p38. One possible mechanism would be the positive regulation of the HDAC class I by ERK1/2, which deacetylates and inhibits the dual-specificity phosphatase 1 (DUSP1; Fig. 7). Acetylated DUSP1 binds with greater affinity to p38, resulting in p38 dephosphorylation. Therefore, the inhibition of DUSP1 by the ERK1/2-HDAC-DUSP1 axis could result in accumulation of more p-p38 in mutant cells (Fig. 7). Our results are consistent with this mechanism, because MEKi reduces p-p38 levels possibly by inhibiting the ERK1/2-HDAC axis and increasing acetylated DUSP1 to target phospho-p38 [17, 21]. Based on previous studies, ERK1/2 was introduced as a negative regulator of p38 *via* inhibition of ASK1-p38 axis (Fig. 2C), however, our data indicate a novel crosstalk between ERK1/2 and p38 in CMs with hyperactive RAF^S257L^. The precise mechanism of the MAPK crosstalk in cardiac myocytes needs further investigation.

### Altered cardiac calcium handling in RAF1^S257L^ cardiomyocytes

One characteristic of maladaptive hypertrophy is an abnormal calcium handling that affects myocardial contractility [11, 34]. In the myocardium, calcium-induced calcium release is essential for excitation-contraction coupling [16]. Depolarization of the plasma membrane through action potential results in L-type calcium channels activation (LTCC) and calcium influx. RYR2 is highly sensitive to small changes in calcium concentration and becomes activated upon local calcium influx, which then facilitates the releases of Ca^2+^ from the sarcoplasmic reticulum into the cytosol. Binding of Ca^2+^ to troponin C causes tropomyosin translocation, which exposes actin filaments for binding to myosin heads, cross-bridge formation and triggers contraction (systole). During diastole, Ca^2+^ is transported into the SR by sarcoplasmic reticulum Ca^2+^-ATPase (SERCA2A).

At the transcriptional level in RAF1^S257L^ CMs, we observed the downregulation of two main regulators of intracellular calcium transients, *SERCA2a* and *LTCC*, as well as changes in the *SERCA2/PLN* ratio. MEKi treatment of RAF1^S257L^ CMs restored the *SERCA2/PLN* ratio and significantly downregulated *NPPB* to levels similar to that of WT CMs. This suggests that SERCA2A/PLN ratio and *NPPB* expression may be under direct or indirect transcriptional control of ERK1/2 (Fig. 7). Previous studies have shown that the RAS-MAPK pathway regulates *SERCA2A*, *PLN*, and *LTCC* expression that is downregulated during hypertrophy and heart failure [6, 27, 28]. Consistently, an increase in BNP, which is known as a HCM biomarker, leads to a decrease in *SERCA2A* expression [70].

In addition to the observed transcriptional alterations of calcium regulators, RAF1^S257L^ CMs also had reduced intracellular Ca^2+^ transients. The influence of hyper-activation of RAS signaling on expression and regulation of SERCA2 and PLN mediated calcium handling and its role in diastolic dysfunction in HCM has been previously reported in mouse models [72]. A lower SERCA2/PLN ratio in RAF1^S257L^ CMs may thereby cause a delay in Ca^2+^ re-entry to the SR *via* inhibition of SERCA by PLN and thus changes the kinetics of calcium transients and consequently decreases the capacity of cardiac contractility [37]. Therefore, a decreased SERCA2a/PLN ratio could be considered as stressor to induce the HCM phenotype [28, 48]. In a similar manner, iPSC-derived CMs with GoF variants in BRAF and MRAS displayed changes in intracellular Ca^2+^ transient [26, 30]. In contrast, a recent study that compared the idiopathic HCM and RAF1^S257L^-associated HCM has shown that CMs from idiopathic cases, but not RAF1^S257L^ CMs, exhibit significant alterations in calcium handling [59].

### Sarcomere disorganization in RAF1^S257L^ cardiomyocytes

RAF1^S257L^ CMs revealed a disorganized sarcomeric structure (Fig. 3A-D). A remarkable and unprecedented finding in the present study is the atypical I-bands, in both RAF1^S257L^ CMs as well as the heart biopsy samples from the corresponding individual with the heterozygous *RAF1^S257L^* variant. We observed this phenotype in several independent experiments, and it was completely re-established upon MEK inhibition.

Immunohistochemistry of the selected region of I-band, the PEVK domains of titin, and Z-line (α-actinin) indicated that two adjacent PEVK regions in RAF1^S257L^ BCTs overlapped (in green) on the Z-line (in red), whereas in WT, the Z-line was clearly surrounded by two distinct and well separated PEVK regions (Fig. 4B). Treatment of BCTs with 0.1 μM MEKi resolved these abnormalities (Fig. 4B and 4C). The I-band segment of titin acts as a molecular spring that develops tension when sarcomeres are stretched, representing a regulatory node that integrates and perhaps coordinates diverse signaling events [33]. The four-and-a-half LIM domain 1 protein (FHL-1) has been shown to bind to titin at the elastic N2B region and to enhance cardiac MAPK signaling by directly interacting as a scaffold protein with RAF1, MEK2, and ERK2 [61]. In our experiments, *FHL1* mRNA expression was up-regulated in RAF1^S257L^ CMs (Fig. S5G). Additionally, we observed that RAF1 was predominantly localized alongside the sarcomeres in RAF1^S257L^ CMs (Fig. 3E). Interestingly, the N2B region of titin has been identified as a substrate of ERK2, and phosphorylation by ERK2 reduces the stiffness of titin [49, 51]. We propose that RAF1^S257L^ hyperactivates the MEK1/2-ERK1/2 pathway, which is most likely localized alongside the sarcomeres, via FHL1, and enhances titin phosphorylation at its N2B region. To what extend these events may result in altered sarcomere distensibility and contributes to the cardiac abnormalities observed with RAF1^S257L^ remains to be determined (Fig. 7, right side).

### RAF1^S257L^ effects on cardiac excitation-contraction coupling

The molecular alterations, structural abnormalities and reduced intracellular Ca^2+^ transients in RAF1^S257L^ CMs expectedly affected contractile behavior of cardiac tissue [15]. Physiological analysis of RAF1^S257L^ BCTs revealed a 1.3-fold higher beating frequency compared to WT BCTs. Treatment of RAF1^S257L^ BCTs with 0.1 and 0.2 μM MEKi decreased the contraction frequency to 0.8- and 0.7-fold of WT BCTs, respectively. The maximum force generation by RAF1^S257L^ BCTs was significantly lower than WT BCTs and inhibiting RAF-MAPK pathway by 0.1 μM MEKi partially restored this value. The RAF1^S257L^ BCTs exhibited reduced contractile tension and 0.1 μM MEKi significantly increased these values to the normal level. Slower contraction might correspond with the reduced Ca^2+^ release shown in Fig. 6F. Furthermore, relaxation would be faster in RAF1^S257L^ than in WT BCTs (Fig. 6F) despite unchanged rates of cytosolic calcium elimination (Fig. 5I).

The link between the RAF1-MAPK signaling pathway and contractile behavior in the myocardium is unclear. One explanation for the altered contractile behavior of RAF1^S257L^ CMs may be the perturbed *MYH6* (α-MYH)-to-*MYH7* (β-MYH) switch due to the aberrantly activated MAPK signaling (Fig. 5B, 7 and S5E). ERK1/2 is known to phosphorylate the cardiac-specific transcription factor GATA4 at S105 and enhances its transcriptional DNA-binding activity. GATA4 is critical for the expression of structural and cardiac hypertrophy response genes, such as *NPPB, MYH7, TNNI3* (troponin I), and *ACTA1* (*α-skeletal Actin*) [2, 4, 10, 25, 36]. We assume that cardiac-specific transcription factors mediate upregulation of myosin heavy chain isoforms by RAF-MAPK. The ATP hydrolyzing capacity of the two myosin heavy chain paralogs are dissimilar; α-MYH has a 3-fold higher ATPase activity and generates more force than β-MYH, which affects the velocity of myofibril shortening and, consequently, contraction [39]. Reduced *MYH6* levels have also been reported in human heart failure [41]. Therefore, we propose that RAF1^S257L^ CMs exhibited higher levels of *MYH7*, leading to less absolute force generation and contractile tension. Additionally, RAF1^S257L^ BCTs need longer time to reach the peak of contraction (Fig. 6E). β-MYH is known as the slower and α-MYH as the faster paralog [20], therefore, RAF1^S257L^ BCTs may need more time to reach the peak of contraction. Alternatively, increased PLN-to-SERCA2 ratio and reduced intracellular calcium transients that may affect intracellular calcium concentrations and cross-bridge cycling kinetics. In addition to the *MYH6-to-MYH7* switch, further factors including SERCA2/PLN ratio, titin phosphorylation by ERK1/2, disorganized sarcomeric structures, and changes in length/shape of the flexible I-band region of titin, might also affect force generation and elastic properties of the myocardium. These changes in the spring elements of titin altering its flexibility may influence timing of contraction and relaxation, where RAF1^S257L^ BCTs need more time for contraction and less time for relaxation (Fig 6E and F). Cardiac contraction-relaxation processes are multifactorial and complementary analysis are required for more clear-cut conclusions.

Collectively, we demonstrated new aspects of RAF1 function in human iPSC-derived CMs, which resemble the observed *in vivo* phenotype from the corresponding individuals, especially changes in the ultra-structure of the sarcomeres. The S257L variant in RAF1 CMs modulates RAF1-dependent signaling networking, fetal gene program, contraction, calcium transients, and the sarcomeric structures. Here, we did not assess the behaviors of all known RAF1 binding partners that may also be critical for cardiac function and contraction, including troponin T, DMPK, ROCK, MYPT, and calcineurin [1, 8, 9, 29, 31, 50, 62, 71]. Therefore, further studies would be supportive to uncover how altered RAF1 signaling impacts these components. We believe future studies with further advanced models of myocardium will uncover more precisely the physiological output of hypertrophic RAF1 variant(s).

## Materials and methods

### Generation, cultivation, and gene-correction of induced pluripotent stem cells

Blood samples and dermal fibroblasts were obtained with the institutional ethics approvals (Justus-Liebig-University Giessen, Germany: AZ258/16, Otto von Guericke University Medical Center Magdeburg: 173/14; University Medical Center Göttingen: 10/9/15) and under informed consent of the parents from two unrelated individuals with Noonan syndrome carrying the heterozygous substitution c.770C>T in exon 7 of *RAF1*.

Primary cells were reprogrammed using either episomal reprogramming vectors Epi5^TM^ [44] (Thermo Scientific #15960) or Sendai virus system Cytotune 2.0 (Thermo Scientific #A16517). Resulting iPSCs were clonally picked and expanded in mTESR (Stemcell Technologies) or StemMACS iPS-Brew (Miltenyi Biotech) to UMGi164-A clone 1 (7B10, here referred to as RAF1^S257L^ line 1) from patient 1 and UMGi102-A clone 17 (isRASb1.17, here referred to as RAF1^S257L^ line 2) from patient 2, respectively, and passaged using Versene (STEMCELL Technologies) at a ratio of 1:3 to 1:6, depending on cell density. Prior to their utilization for experiments, clonal iPSCs were subjected to detailed characterization including Sanger sequencing to confirm the presence of the variant, iPSC morphology, assessment of expression of pluripotency markers by RT-PCR, immunofluorescence staining and flow cytometry, and chromosomal integrity. Furthermore, elimination of persisting reprogramming factors was confirmed by PCR and RT-PCR, respectively. Three unrelated wild-type (WT) iPSC lines, UMGi163-A clone 1 (ipWT16.1, here referred to as WT1) [22], UMGi014-C clone 14 (isWT1.14, here referred to as WT2) and UMGi020-B clone 22 (isWT7.22, here referred to as WT3) [58]. A CRISPR-corrected isogenic iPSC line was used as control. Genetic correction of the RAF^S257L^ variant (c.770 C>T, heterozygous) in the iPSCs from patient 2 was performed using ribonucleoprotein (RNP)-based CRISPR/Cas9 by targeting exon 7 of the *RAF1* gene, as previously described [24]. The guide RNA target sequence was (PAM in bold): 5’-TGGATGTCAACCTCTGCCTC <u>TGG</u>-3’. For homology-directed repair, a single-stranded oligonucleotide with 45-bp homology arms was used. After picking clones, successful gene-editing was identified by Sanger sequencing and the CRISPR-corrected isogenic cell line UMGi102-A-1 clone 9 (isRASb1-corr.9) underwent the same detailed characterization as mentioned above.

#### Human iPSC culture

Before initiation of cardiomyogenic differentiation, undifferentiated iPSCs were cultured as feeder-free monolayers for up to 15 passages in murine embryonic feeder cell-conditioned medium (CCM^+^) consisting of DMEM F12+Glutamax, 15% Knock-out Serum Replacement, 1% non-essential amino acids (all Gibco), 100 μM β-mercaptoethanol (Sigma), and 100 ng/mL bFGF (PeproTech; CCM^+^/100). Cells were passaged every three to four days by dissociation with Accutase and seeded onto Geltrex-coated (0.5%, Life Technologies) plasticware at a density of 5×10^4^ cells per cm^2^ in CCM^+^, containing the ROCK inhibitor Y-27632 (10 μM, Selleckchem, #S1049). The ROCK inhibitor was eliminated from the CCM^+^ on the following days.

### Tri-lineage differentiation of human iPSCs

To induce differentiation of human iPSCs into all three germ layers, human iPSC-colonies were detached from feeder layers using 0.4% (w/v) type IV collagenase and resuspended in differentiation medium consisting of IMDM + GlutaMAX supplemented with 20% (v/v) fetal calf serum, 1 mL L-Glutamine, 0.1 mM 2-mercaptoethanol, and 1% non-essential amino acid stock (all Thermo Scientific). Colonies were maintained for 7 days in suspension culture on 1% (w/v) agarose/IMDM coated 12-well plates to form 3D embryoid bodies (EBs). Subsequently, about 15-20 EBs were plated on 6-well plates coated with 0.1% (w/v) gelatine. After 24 days, EBs were harvested for qRT-PCR and replated for immunofluorescence (IF) analyses, respectively.

### Karyotype analysis

After treatment of undifferentiated human iPSCs with a final concentration of 0.1 μg/mL Colcemid (Thermo Scientific) for 2 h, cells were detached with trypsin/EDTA (0.05/0.02%, Biochrom). After centrifugation, the pellet was resuspended in hypotonic solution (0.32% KCl with 0.2% (v/v) fetal calf serum) and incubated for 15 min at 37 °C. Cells were fixed in ice-cold methanol/acetic acid (3:1). G-banding was performed according to Seabright protocol[60]. Karyograms were imaged using the IKAROS software of MetaSystems (Altluβheim, Germany). The chromosome arrangement was investigated as previously described [23].

### Cardiac differentiation of human iPSCs

Cardiac differentiation was performed in 3D suspension culture after aggregate formation in agarose microwells modified from established protocol [5, 35]. For differentiation in 3D suspension culture, agarose microwells (AMW) were generated from AggreWellTM400Ex plates (Stem Cell Technologies, #27840) containing 4700 microslots per AMW in a 6-well format [5]. For each AMW, 5×10^6^ undifferentiated human iPSCs were seeded in 3 mL CCM^+^/100 supplemented with 10 μM ROCK inhibitor Y-27632 (Selleckchem, #S1049). iPSCs formed uniform EBs during the initial 24 h on AMW, and were harvested and transferred to suspension culture in 15-cm dishes and placed on an orbital shaker at 60 rpm. Suspension EBs were cultivated for further 3 days in CCM^+^/100 before the start of cardiac differentiation. Differentiation was induced with the exchange of medium to RPMI 1640 supplemented with 1% (v/v) B27 without insulin (RB^-^, Thermo Scientific, #A18956-01). The GSK-3 inhibitor CHIR99021 (Selleckchem, #S1263) was added at 4 - 6 μM (depending on the iPSC line) for the first 24 h of differentiation, thereafter differentiations were kept in RB^-^ without small molecule inhibitors until day 3 (d3) before the WNT inhibitor IWR-1 (4 μM, Sigma, #I0161) was added for 48 h. The first contracting EBs were observed between d5 and d7. From d7-d10, EBs were cultivated with RPMI 1640 supplemented with 1% (v/v) B27 with insulin (RB^+^, Thermo Scientific, #17504-044). Afterwards, a metabolic selection was performed for 10d (until d20) to eliminate all non-CM in RPMI minus glucose (Thermo Scientific, #11879-020) supplemented with human albumin (Sigma, #A0237), sodium DL-lactate (Sigma, #L4263), and L-ascorbic acid-2-phosphate (Sigma, #A8960) [64]. From d20 - d40, CBs were kept in RB^+^ medium. Depending on the experimental design, inhibition of MEK was started on d12 of differentiation by supplementing the medium with 0.1 and 0.2 μM PD0325901 (Sigma, #PZ0162). Medium was exchanged every two days until d20. Flow cytometric assessment of CM content was done on d20 by staining against cardiac troponin T (Life technologies) after dissociation of aggregates with Stemdiff CM dissociation kit (STEMCELL Technologies) following the manufacturer’s protocol after adding Accutase to the enzyme mix (1:2). Aggregates with a purity of >95% were termed cardiac bodies (CBs) and further cultivated.

### Bioartificial cardiac tissue (BCT) preparation, culture, and measurements

On day 21, non-MEK-inhibited CBs were dissociated as described above and a previously described protocol to generate myocardial tissue was employed [32]. In brief, a mixture of rat tail collagen type I (Cultrex) and Matrigel™ (Life Technologies) was mixed with CMs and gamma-irradiated human foreskin fibroblasts (HFFs) and poured into silicone molds (5×5×10 mm, w/d/h) with two horizontal titanium rods that serve as suspensions for the developing tissue at a distance of 6 mm. Resulting BCTs with a volume of 250 μL each containing 10^6^ CMs, 10^5^ HFFs, collagen type I (230 μg), and 10% Matrigel. BCTs were stretched by 200 μm every fourth day to support tissue maturation, and tissue development was documented by daily microscopic observation. Beating frequencies, active contraction, and passive forces were determined in a custom-made bioreactor system at 200 μm increments until the maximum contraction force was reached (L_max_). Contractile and passive tension was calculated by dividing the force by the cross-sectional area of each BCT (mN/mm^2^).

### Reverse transcriptase polymerase chain reaction

Cells were lysed using TRIzol™ (Ambion, Thermo Scientific, Germany) and total RNA was extracted via phenol-chloroform extraction. Remaining genomic DNA contaminations were removed using the DNA-free™ DNA Removal Kit (Ambion, Thermo Scientific, Germany). DNase-treated RNA was transcribed into complementary DNA (cDNA) using the ImProm-II™ reverse transcription system and oligo-dT as primer (Promega, Germany). Quantitative real-time reverse transcriptase polymerase chain reaction (qPCR) was performed using SYBR Green (Thermo Scientific, #4309155 Germany). Primer sequences are listed in Supplementary Table S1. The 2^-ΔΔCt^ method was employed for estimating the relative mRNA expression levels and 2^-ΔCt^ for mRNA levels. Among six different housekeeping genes that we tested, *HPRT1* showed the lowest variation among different cell lines and conditions. Therefore, *HPRT1* was used for normalization in our qPCR analysis.

### Flow cytometry

For flow cytometric analysis, single-cell suspensions of undifferentiated human iPSCs were obtained with Accutase and cells were washed with ice-cold phosphate-buffered saline (PBS)^-/-^. CBs were dissociated into single cells by incubation with Versene (EDTA-Solution, Thermo Scientific, #15040066) for 10 min in a Thermomixer at 37°C. Thereafter, TrypLE (Thermo Scientific, #A1285901) was added and samples were incubated for additional 10 min at 37 °C and 1200 rpm until the cellular aggregates have dissolved. Both cell types were fixed in 4% formaldehyde (Carl Roth, #P087.1) for 10 min on ice and permeabilized with 90% ice-cold methanol for 20 min followed by a blocking step with 1.5% BSA and 2.5% goat or donkey serum diluted in PBS for 1 h, at 4 °C. Cells were then stained with primary antibodies.

### Immunoblotting

To extract the total protein, CBs were washed with PBS^-/-^ prior to cell lysis (lysis buffer: 50 mM Tris-HCl pH 7.5; 100 mM NaCl, 2 mM MgCl_2_; 1% Triton X100; 10% glycerol; 20 mM beta-glycerol phosphate; 1 mM orthoNa_3_VO_4_; and EDTA-free protease inhibitor (Roche, Germany, #11873580001)). To further disrupt the cellular aggregates, a sonicator with 70% power was used prior to incubation in a rotor at 4 °C for 30 min. Protein concentrations were determined using the Bradford assay (Bio-Rad, #5000201). Equal amounts of cell lysates (10 - 50 μg) were subjected to sodium dodecyl sulfate polyacrylamide gel electrophoresis (SDS-PAGE). Following electrophoresis, proteins were transferred to a nitrocellulose membrane by electroblotting and probed with primary antibodies overnight at 4°C. All antibodies were diluted in Intercept® (TBS) Blocking Buffer (LI-COR #927-60001) mixed in a 1:3 ratio with Tris-buffered saline containing 0.02% Tween 20. The antibodies from Santa Cruz were diluted 1:200 while the remaining antibodies were diluted 1:1000 (see Table S2). Immunoblots were developed with LI-COR Odyssey FX (LI-COR) and quantification of signals was performed by densitometry of scanned signals with the aid of Image Studio (version 5.2, LI-COR).

### Immunocytochemistry

Immunostaining was performed as described previously [43]. The procedure for the single-cell suspensions of CBs was described in the flow cytometry section. Briefly, cells were washed twice with ice-cold PBS containing magnesium/calcium and fixed with 4% formaldehyde (Carl Roth, #P087.1) for 20 min at room temperature. To permeabilize cell membranes, cells were incubated in 0.25% Triton X-100/PBS for 5 min. Blocking was performed with 3% bovine serum albumin (Thermo Scientific, # 26140079) and 2% goat serum diluted in PBS containing 0.25% Triton X-100 for 1 h at room temperature. Incubation with primary antibodies was performed overnight. Cells were washed three times for 10 min with PBS and incubated with secondary antibodies for 2 h at room temperature. Slides were washed three times and ProLong® Gold Antifade mounting reagent containing DAPI (4’,6-diamidino-2-phenylindole) (Thermo Scientific, #P10144) was applied to mount the coverslips.

### Immunohistochemistry

From formalin fixed tissue, 3 μm sections were stained with H&E and Masson trichrome (Trichrome II Blue staining kit at Nexus special stainer; Roche). Immunohistochemical analysis was performed on cryosections and paraffin sections using a Bench Mark XT automatic staining platform (Ventana, Heidelberg, Germany) with the primary antibodies. The sections were examined using a Nikon Eclipse 80i equipped with a DS-Fi1 camera.

A myectomy was performed on the patient one with RAF mutation at the age of 5 years. After formalin-fixation of the surgically removed tissue, 3μm sections were stained with hematoxylin and eosin. Immunohistochemical analysis was performed on cryosections and paraffin sections using a Bench Mark XT automatic staining platform (Ventana, Heidelberg, Germany), using the primary antibodies listed in the supplementary table S2. The sections were examined using a Nikon Eclipse 80i equipped with a DS-Fi1 camera.

Immunohistochemistry of BCT samples was carried out using BCTs embedded and snap-frozen in Tissue-Tek® OCT resin (Sakura Finetek, #4583). The cryo-blocks were sliced into 8μM thick sections. Sections were fixed in 4% paraformaldehyde (Carl Roth, #P087.1) in 0.1 mol/L sodium phosphate buffer pH 7.4 for 10 min. Washing was performed in PBS and 0.2% saponin/PBS. After blocking with 10% normal goat serum (NGS) in 0.2% saponin/PBS for 1 h, primary antibodies were incubated over night at 4°C. Secondary antibodies were incubated for 3 h at room temperature in the dark. The sections were mounted with ProLong® Gold Antifade Mountant, containing ProLDAPI (#P-36935, Invitrogen). Slides were analyzed with a ZEISS Airyscan LSM 880 confocal microscope (Center for Advanced Imaging, Heinrich Heine University, Düsseldorf, Germany). Pictures were taken with 63x objective and analyzed using ZEN 3.2 (blue edition) by Carl Zeiss AG.

### Immunoprecipitation

For immunoprecipitation of RAF1, beads were initially incubated with an anti-RAF1 antibody and a non-specific anti-IgG for the negative control, respectively, and washed afterwards five times with immunoprecipitation buffer (20 mM Tris/HCl pH 7.4, 150 mM NaCl, 5 mM MgCl_2_, 10 mM β-glycerophosphate, 0.5 mM Na_3_VO_4_, 10% glycerol, and EDTA-free protease inhibitor). Afterwards, iPSCs cells were lysed in lysis buffer (immunoprecipitation buffer with 0.5% NP-40) by 30 min rotation in a reaction tube at 4°C. Immunoprecipitation from total cell lysates was carried out for 1 h, at 4°C. Beads were washed five times with immunoprecipitation buffer, and eluted proteins were heated in SDS/Laemmli sample buffer at 95°C for 10 min before subjected to immunoblotting.

### Analysis of cardiomyocyte cell surface

Analysis of CM cell surface was conducted using a Java-based open-source image processing software ImageJ. Software pixel length was calibrated to the μm scale bar on the used confocal images. After background elimination, the Region of Interest-tool (ROI) was used to mark the borders of single cells. For each condition, multiple confocal images were evaluated in order to evaluate a sufficient number of images. Finally, the cell area was extracted and compared between the conditions.

### Transmission electron microscopy (TEM)

Small cardiac tissue samples were fixed with 6% glutaraldehyde/0.4 M phosphate buffered saline (PBS) and processed with a Leica EM TP tissue processor. Cardiac bodies were fixed with 3% glutaraldehyde/0.1 M Cacodylate buffer. The cell pellets were processed by hand according to the automated tissue processor. For electron microscopy of the small cardiac tissue samples, ultrathin sections were contrasted with 3% lead citrate trihydrate with a Leica EM AC20 (Ultrastain kit II) and were examined using a ZEISS EM 109 transmission electron microscope equipped with a Slowscan-2K-CCD-digital camera (2K-wide-angle Sharp: eye), while CBs were imaged using a Hitachi H-7100.

### Measurement of Ca^2+^ cycling in cardiac bodies

iPSC-derived CBs were dissociated and grown on gelatin coated cover slips for up to 7 d before loading with the fluorescent Ca^2+^ indicator Fura-2 by adding 1 μg Fura-2-AM/mL cell medium. After a 15 min incubation at 37°C, cells were washed in pre-warmed medium (37°C). A dual excitation (340 nm and 380 nm) fluorescence imaging recording system was used to measure Ca^2+^ transients of paced (0.5 Hz) and spontaneously beating cells (HyperSwitch Myocyte System, IonOptix Corp., Milton, MA, USA). Data were acquired as the ratio of measurements at 340 and 380 nm and analyzed using IonWizard software (Version 6.4, Ion Optix Corp).

## Acknowledgements

We are grateful to Dr. Ehsan Amin and Prof. Dr. Thomas Wieland for helpful advices, and stimulating discussions. We are grateful to the CFEM (Core Facility Electron Microscopy) of the medical faculty of the Heinrich Heine University Düsseldorf. We gratefully thank the entire team from the Stem Cell Unit, University Medical Center Göttingen, for excellent technical assistance in iPSC generation and characterization. The authors thank Louisa Habich, Sarah Nourmohammadi, Hannah Schlierbach, David Skvorc, Marion Möckel and Kerstin Leib for excellent technical assistance in iPSC differentiation, CM selection and BCT production. We thank and give credit to “BioRender.com” which was used for design and creation of Fig. 1A.

## Ethical Approval and Consent to participate

Blood samples and dermal fibroblasts were obtained by skin biopsy from two unrelated female patients with Noonan syndrome carrying the heterozygous substitution c.770C>T in exon 7 of RAF1 under protocols concerning research with biomaterials approved by the institution’s ethics committees (Justus-Liebig-University Giessen, Germany: AZ258/16; Otto von Guericke University Medical Center Magdeburg: 173/14; University Medical Center Göttingen: 10/9/15). Research with biomaterials was approved by the ethical committee of the Justus Liebig University of Giessen. Written informed consent was given by both parents of both individuals. One human iPSC line that was used in this study as a control, was generated from the human foreskin fibroblast-1 (HFF-1) cell line (ATCC).

## Authors’ contributions

Conception, design, and writing: MRA, SNR and FB

Development of methodology: SNR, FB, MB, JD, FH, MK, AB, RA, GK, LC

Acquisition of data, analysis and interpretation of data: SNR, FB, AVB, MB, FH, JD, FF, SK, AB, AVK, ASR, AS, MT, GK

Administrative, technical, or material support: MRA, GK, MZ, MK, AG, MJW, AS, ASR, AH, RPP, JS, LC

Study supervision and coordination: MRA, GK, SNR and MZ

## Abbreviations

AC: adenylyl cyclase
ACTN1: actinin alpha 1
APF: alpha fetoprotein
α-SMA: alpha smooth muscle actin
BNP: brain natriuretic peptide
CACNA1C: calcium voltage-gated channel subunit alpha1 C
CB: cardiac body
CT: cardiac tissue
CM: cardiomyocytes
cTNT: cardiac troponin T
DUSP1: dual-specificity phosphatase
EB: embryoid body
EBNA1: Epstein-Barr nuclear antigen 1
EM: electron microscopy
ERK: extracellular regulated kinase
FOXOA2: forkhead box A2
GSK3β: glycogen synthase kinase 3 beta
HCM: hypertrophic cardiomyopathy
HDAC: histone deacetylase
HPRT1: hypoxanthine-guanine phosphoribosyltransferase
LTCC: L-type calcium channels
MAPK: mitogen-activated protein kinase
MEF2: myocyte enhancer factor 2
MEK: MAP/ERK kinase
MEKi: MAP/ERK kinase inhibitor
MYH: myosin heavy chain
MYL: myosin light chain 7
iPSC: induced pluripotent stem cells
NFAT: nuclear factor of activated T-cells
NC: nocodazole
NKX2.5: NK2 homeobox 5
NPPB: natriuretic peptide B
NS: Noonan syndrome
OCT-4: octamer-binding transcription factor 4
PE: phenylephrine
p-H3: phosphohistone 3
PLN: phospholamban
RAF: rapidly accelerated fibrosarcoma
RT-PCR: reverse transcriptase polymerase chain
RyR2: ryanodine receptor type-2
SERCA2A: sarco/endoplasmic reticulum Ca^2+^-ATPase
SOX2: SRY-box transcription factor 2
TNNC1: troponin C1
TNNI3: troponin I3, cardiac type
TRP53: transformation related protein 53
TUBB3: tubulin beta 3 class III
WT: wild-type.

## References

1. Broustas CG, Grammatikakis N, Eto M, Dent P, Brautigan DL, Kasid U (2001) Phosphorylation of the Myosin-binding Subunit of Myosin Phosphatase by Raf-1 and Inhibition of Phosphatase Activity. J Biol Chem 277:3053–3059 doi:10.1074/jbc.M106343200

2. Bueno OF, De Windt LJ, Tymitz KM, Witt SA, Kimball TR, Klevitsky R, Hewett TE, Jones SP, Lefer DJ, Peng CF, Kitsis RN, Molkentin JD (2000) The MEK1-ERK1/2 signaling pathway promotes compensated cardiac hypertrophy in transgenic mice. EMBO J 19:6341–50 doi:10.1093/emboj/19.23.6341

3. Calcagni G, Adorisio R, Martinelli S, Grutter G, Baban A, Versacci P, Digilio MC, Drago F, Gelb BD, Tartaglia M, Marino B (2018) Clinical Presentation and Natural History of Hypertrophic Cardiomyopathy in RASopathies. Heart Fail Clin 14:225–235 doi:10.1016/j.hfc.2017.12.005

4. Charron F, Paradis P, Bronchain O, Nemer G, Nemer M (1999) Cooperative interaction between GATA-4 and GATA-6 regulates myocardial gene expression. Mol Cell Biol 19:4355–65 doi:10.1128/mcb.19.6.4355

5. Dahlmann J, Kensah G, Kempf H, Skvorc D, Gawol A, Elliott DA, Drager G, Zweigerdt R, Martin U, Gruh I (2013) The use of agarose microwells for scalable embryoid body formation and cardiac differentiation of human and murine pluripotent stem cells. Biomaterials 34:2463–71 doi:10.1016/j.biomaterials.2012.12.024

6. Daub M, Jockel J, Quack T, Weber CK, Schmitz F, Rapp UR, Wittinghofer A, Block C (1998) The RafC1 cysteine-rich domain contains multiple distinct regulatory epitopes which control Ras-dependent Raf activation. Mol Cell Biol 18:6698–710 doi:10.1128/mcb.18.11.6698

7. Del Re DP, Yang Y, Nakano N, Cho J, Zhai P, Yamamoto T, Zhang N, Yabuta N, Nojima H, Pan D, Sadoshima J (2012) Yes-associated Protein Isoform 1 (Yap1) Promotes Cardiomyocyte Survival and Growth to Protect against Myocardial Ischemic Injury. J Biol Chem 288:3977–3988 doi:10.1074/jbc.M112.436311

8. Desideri E, Cavallo AL, Baccarini M (2015) Alike but Different: RAF Paralogs and Their Signaling Outputs. Cell 161:967–970 doi:10.1016/j.cell.2015.04.045

9. Dhandapany PS, Fabris F, Tonk R, Illaste A, Karakikes I, Sorourian M, Sheng J, Hajjar RJ, Tartaglia M, Sobie EA, Lebeche D, Gelb BD (2011) Cyclosporine attenuates cardiomyocyte hypertrophy induced by RAF1 mutants in Noonan and LEOPARD syndromes. J Mol Cell Cardiol 51:4–15 doi:10.1016/j.yjmcc.2011.03.001

10. Dirkx E, da Costa Martins PA, De Windt LJ (2013) Regulation of fetal gene expression in heart failure. Biochim Biophys Acta 1832:2414–24 doi:10.1016/j.bbadis.2013.07.023

11. Dorn GW, 2nd, Force T (2005) Protein kinase cascades in the regulation of cardiac hypertrophy. J Clin Invest 115:527–37 doi:10.1172/jci24178

12. Dougherty MK, Müller J, Ritt DA, Zhou M, Zhou XZ, Copeland TD, Conrads TP, Veenstra TD, Lu KP, Morrison DK (2005) Regulation of Raf-1 by direct feedback phosphorylation. Mol Cell 17:215–24 doi:10.1016/j.molcel.2004.11.055

13. Doyle MJ, Lohr JL, Chapman CS, Koyano-Nakagawa N, Garry MG, Garry DJ (2015) Human Induced Pluripotent Stem Cell-Derived Cardiomyocytes as a Model for Heart Development and Congenital Heart Disease. Stem Cell Rev 11:710–27 doi:10.1007/s12015-015-9596-6

14. Dumaz N, Marais R (2003) Protein kinase A blocks Raf-1 activity by stimulating 14-3-3 binding and blocking Raf-1 interaction with Ras. J Biol Chem 278:29819–29823 doi:10.1074/jbc.C300182200

15. Eisner DA, Caldwell JL, Kistamas K, Trafford AW (2017) Calcium and Excitation-Contraction Coupling in the Heart. Circ Res 121:181–195 doi:10.1161/circresaha.117.310230

16. Fabiato A (1983) Calcium-induced release of calcium from the cardiac sarcoplasmic reticulum. Am J Physiol 245:C1–14 doi:10.1152/ajpcell.1983.245.1.C1

17. Ferguson BS, Harrison BC, Jeong MY, Reid BG, Wempe MF, Wagner FF, Holson EB, McKinsey TA (2013) Signal-dependent repression of DUSP5 by class I HDACs controls nuclear ERK activity and cardiomyocyte hypertrophy. Proc Natl Acad Sci U S A 110:9806–11 doi:10.1073/pnas.1301509110

18. Gallo S, Vitacolonna A, Bonzano A, Comoglio P, Crepaldi T (2019) ERK: A Key Player in the Pathophysiology of Cardiac Hypertrophy. Int J Mol Sci 20 doi:10.3390/ijms20092164

19. Gelb BD, Roberts AE, Tartaglia M (2015) Cardiomyopathies in Noonan syndrome and the other RASopathies. Prog Pediatr Cardiol 39:13–19 doi:10.1016/j.ppedcard.2015.01.002

20. Gupta MP (2007) Factors controlling cardiac myosin-isoform shift during hypertrophy and heart failure. J Mol Cell Cardiol 43:388–403 doi:10.1016/j.yjmcc.2007.07.045

21. Habibian J, Ferguson BS (2018) The Crosstalk between Acetylation and Phosphorylation: Emerging New Roles for HDAC Inhibitors in the Heart. Int J Mol Sci 20 doi:10.3390/ijms20010102

22. Haghighi F, Dahlmann J, Nakhaei-Rad S, Lang A, Kutschka I, Zenker M, Kensah G, Piekorz RP, Ahmadian MR (2018) bFGF-mediated pluripotency maintenance in human induced pluripotent stem cells is associated with NRAS-MAPK signaling. Cell Commun Signal 16:96 doi:10.1186/s12964-018-0307-1

23. Hamta A, Adamovic T, Samuelson E, Helou K, Behboudi A, Levan G (2006) Chromosome ideograms of the laboratory rat (Rattus norvegicus) based on high-resolution banding, and anchoring of the cytogenetic map to the DNA sequence by FISH in sample chromosomes. Cytogenet Genome Res 115:158–68 doi:10.1159/000095237

24. Hanses U, Kleinsorge M, Roos L, Yigit G, Li Y, Barbarics B, El-Battrawy I, Lan H, Tiburcy M, Hindmarsh R (2020) Intronic CRISPR Repair in a Preclinical Model of Noonan Syndrome–Associated Cardiomyopathy. Circulation 142:1059–1076 doi:10.1161/CIRCULATIONAHA.119.044794

25. He A, Kong SW, Ma Q, Pu WT (2011) Co-occupancy by multiple cardiac transcription factors identifies transcriptional enhancers active in heart. Proc Natl Acad Sci U S A 108:5632–7 doi:10.1073/pnas.1016959108

26. Higgins EM, Bos JM, Dotzler SM, Kim CJ, Ackerman MJ (2019) MRAS Variants Cause Cardiomyocyte Hypertrophy in Patient-Specific iPSC-Derived Cardiomyocytes: Additional Evidence for MRAS as a Definitive Noonan Syndrome-Susceptibility Gene. Circ Genom Precis Med doi:10.1161/circgen.119.002648

27. Ho PD, Fan JS, Hayes NL, Saada N, Palade PT, Glembotski CC, McDonough PM (2001) Ras reduces L-type calcium channel current in cardiac myocytes. Corrective effects of L-channels and SERCA2 on [Ca(2+)](i) regulation and cell morphology. Circ Res 88:63–9 doi:10.1161/01.res.88.1.63

28. Huang H, Joseph LC, Gurin MI, Thorp EB, Morrow JP (2014) Extracellular signal-regulated kinase activation during cardiac hypertrophy reduces sarcoplasmic/endoplasmic reticulum calcium ATPase 2 (SERCA2) transcription. J Mol Cell Cardiol 75:58–63 doi:10.1016/j.yjmcc.2014.06.018

29. Jaffre F, Miller CL, Schanzer A, Evans T, Roberts AE, Hahn A, Kontaridis MI (2019) Inducible Pluripotent Stem Cell-Derived Cardiomyocytes Reveal Aberrant Extracellular Regulated Kinase 5 and Mitogen-Activated Protein Kinase Kinase 1/2 Signaling Concomitantly Promote Hypertrophic Cardiomyopathy in RAF1-Associated Noonan Syndrome. Circulation 140:207–224 doi:10.1161/circulationaha.118.037227

30. Josowitz R, Mulero-Navarro S, Rodriguez NA, Falce C, Cohen N, Ullian EM, Weiss LA, Rauen KA, Sobie EA, Gelb BD (2016) Autonomous and Non-autonomous Defects Underlie Hypertrophic Cardiomyopathy in BRAF-Mutant hiPSC-Derived Cardiomyocytes. Stem Cell Reports 7:355–369 doi:10.1016/j.stemcr.2016.07.018

31. Kaliman P, Llagostera E (2008) Myotonic dystrophy protein kinase (DMPK) and its role in the pathogenesis of myotonic dystrophy 1. Cell Signal 20:1935–1941 doi:10.1016/j.cellsig.2008.05.005

32. Kensah G, Roa Lara A, Dahlmann J, Zweigerdt R, Schwanke K, Hegermann J, Skvorc D, Gawol A, Azizian A, Wagner S, Maier LS, Krause A, Drager G, Ochs M, Haverich A, Gruh I, Martin U (2013) Murine and human pluripotent stem cell-derived cardiac bodies form contractile myocardial tissue in vitro. Eur Heart J 34:1134–46 doi:10.1093/eurheartj/ehs349

33. Krüger M, Linke WA (2011) The Giant Protein Titin: A Regulatory Node That Integrates Myocyte Signaling Pathways. J Biol Chem 286:9905–9912 doi:10.1074/jbc.R110.173260

34. Lan F, Lee AS, Liang P, Sanchez-Freire V, Nguyen PK, Wang L, Han L, Yen M, Wang Y, Sun N, Abilez OJ, Hu S, Ebert AD, Navarrete EG, Simmons CS, Wheeler M, Pruitt B, Lewis R, Yamaguchi Y, Ashley EA, Bers DM, Robbins RC, Longaker MT, Wu JC (2013) Abnormal calcium handling properties underlie familial hypertrophic cardiomyopathy pathology in patient-specific induced pluripotent stem cells. Cell Stem Cell 12:101–13 doi:10.1016/j.stem.2012.10.010

35. Lian X, Zhang J, Azarin SM, Zhu K, Hazeltine LB, Bao X, Hsiao C, Kamp TJ, Palecek SP (2013) Directed cardiomyocyte differentiation from human pluripotent stem cells by modulating Wnt/beta-catenin signaling under fully defined conditions. Nat Protoc 8:162–75 doi:10.1038/nprot.2012.150

36. Liu YL, Huang CC, Chang CC, Chou CY, Lin SY, Wang IK, Hsieh DJ, Jong GP, Huang CY, Wang CM (2015) Hyperphosphate-Induced Myocardial Hypertrophy through the GATA-4/NFAT-3 Signaling Pathway Is Attenuated by ERK Inhibitor Treatment. Cardiorenal Med 5:79–88 doi:10.1159/000371454

37. MacLennan DH, Kranias EG (2003) Phospholamban: a crucial regulator of cardiac contractility. Nat Rev Mol Cell Biol 4:566–77 doi:10.1038/nrm1151

38. Maron BJ, Maron MS (2013) Hypertrophic cardiomyopathy. The Lancet 381:242–255 doi:10.1016/s0140-6736(12)60397-3

39. Miyata S, Minobe W, Bristow MR, Leinwand LA (2000) Myosin heavy chain isoform expression in the failing and nonfailing human heart. Circ Res 86:386–90 doi:10.1161/01.res.86.4.386

40. Mosqueira D, Mannhardt I, Bhagwan JR, Lis-Slimak K, Katili P, Scott E, Hassan M, Prondzynski M, Harmer SC, Tinker A, Smith JGW, Carrier L, Williams PM, Gaffney D, Eschenhagen T, Hansen A, Denning C (2018) CRISPR/Cas9 editing in human pluripotent stem cell-cardiomyocytes highlights arrhythmias, hypocontractility, and energy depletion as potential therapeutic targets for hypertrophic cardiomyopathy. Eur Heart J 39:3879–3892 doi:10.1093/eurheartj/ehy249

41. Nakao K, Minobe W, Roden R, Bristow MR, Leinwand LA (1997) Myosin heavy chain gene expression in human heart failure. J Clin Invest 100:2362–70 doi:10.1172/jci119776

42. Nakhaei-Rad S, Bahrami AR, Mirahmadi M, Matin MM (2012) New windows to enhance direct reprogramming of somatic cells towards induced pluripotent stem cells. Biochem Cell Biol 90:115–23 doi:10.1139/o11-064

43. Nakhaei-Rad S, Nakhaeizadeh H, Kordes C, Cirstea IC, Schmick M, Dvorsky R, Bastiaens PI, Haussinger D, Ahmadian MR (2015) The Function of Embryonic Stem Cell-expressed RAS (ERAS), a Unique RAS Family Member, Correlates with Its Additional Motifs and Its Structural Properties. J Biol Chem 290:15 892-903 doi:10.1074/jbc.M115.640607

44. Okita K, Matsumura Y, Sato Y, Okada A, Morizane A, Okamoto S, Hong H, Nakagawa M, Tanabe K, Tezuka K, Shibata T, Kunisada T, Takahashi M, Takahashi J, Saji H, Yamanaka S (2011) A more efficient method to generate integration-free human iPS cells. Nat Methods 8:409–12 doi:10.1038/nmeth.1591

45. Olivotto I, d’Amati G, Basso C, Van Rossum A, Patten M, Emdin M, Pinto Y, Tomberli B, Camici PG, Michels M (2015) Defining phenotypes and disease progression in sarcomeric cardiomyopathies: contemporary role of clinical investigations. Cardiovasc Res 105:409–423 doi:10.1093/cvr/cvv024

46. Ovchinnikova E, Hoes M, Ustyantsev K, Bomer N, de Jong TV, van der Mei H, Berezikov E, van der Meer P (2018) Modeling Human Cardiac Hypertrophy in Stem Cell-Derived Cardiomyocytes. Stem Cell Reports 10:794–807 doi:10.1016/j.stemcr.2018.01.016

47. Pandit B, Sarkozy A, Pennacchio LA, Carta C, Oishi K, Martinelli S, Pogna EA, Schackwitz W, Ustaszewska A, Landstrom A, Bos JM, Ommen SR, Esposito G, Lepri F, Faul C, Mundel P, López Siguero JP, Tenconi R, Selicorni A, Rossi C, Mazzanti L, Torrente I, Marino B, Digilio MC, Zampino G, Ackerman MJ, Dallapiccola B, Tartaglia M, Gelb BD (2007) Gain-of-function RAF1 mutations cause Noonan and LEOPARD syndromes with hypertrophic cardiomyopathy. Nat Genet 39:1007–1012 doi:10.1038/ng2073

48. Periasamy M, Bhupathy P, Babu GJ (2007) Regulation of sarcoplasmic reticulum Ca2+ ATPase pump expression and its relevance to cardiac muscle physiology and pathology. Cardiovasc Res 77:265–273 doi:10.1093/cvr/cvm056

49. Perkin J, Slater R, Del Favero G, Lanzicher T, Hidalgo C, Anderson B, Smith JE, 3rd, Sbaizero O, Labeit S, Granzier H (2015) Phosphorylating Titin’s Cardiac N2B Element by ERK2 or CaMKIIδ Lowers the Single Molecule and Cardiac Muscle Force. Biophys J 109:2592–2601 doi:10.1016/j.bpj.2015.11.002

50. Pfleiderer P, Sumandea MP, Rybin VO, Wang C, Steinberg SF (2009) Raf-1: a novel cardiac troponin T kinase. J Muscle Res Cell Motil 30:67–72 doi:10.1007/s10974-009-9176-y

51. Raskin A, Lange S, Banares K, Lyon RC, Zieseniss A, Lee LK, Yamazaki KG, Granzier HL, Gregorio CC, McCulloch AD, Omens JH, Sheikh F (2012) A novel mechanism involving four-and-a-half LIM domain protein-1 and extracellular signal-regulated kinase-2 regulates titin phosphorylation and mechanics. J Biol Chem 287:29273–84 doi:10.1074/jbc.M112.372839

52. Rauen KA (2013) The RASopathies. Annu Rev Genomics Hum Genet 14:355–369 doi:10.1146/annurev-genom-091212-153523

53. Razzaque MA, Nishizawa T, Komoike Y, Yagi H, Furutani M, Amo R, Kamisago M, Momma K, Katayama H, Nakagawa M, Fujiwara Y, Matsushima M, Mizuno K, Tokuyama M, Hirota H, Muneuchi J, Higashinakagawa T, Matsuoka R (2007) Germline gain-of-function mutations in RAF1 cause Noonan syndrome. Nat Genet 39:1013–1017 doi:10.1038/ng2078

54. Rezaei Adariani S, Buchholzer M, Akbarzadeh M, Nakhaei-Rad S, Dvorsky R, Ahmadian MR (2018) Structural snapshots of RAF kinase interactions. Biochem Soc Trans 46:1393–1406 doi:10.1042/bst20170528

55. Roberts AE, Allanson JE, Tartaglia M, Gelb BD (2013) Noonan syndrome. The Lancet 381:333–342 doi:10.1016/S0140-6736(12)61023-X

56. Romano D, Nguyen LK, Matallanas D, Halasz M, Doherty C, Kholodenko BN, Kolch W (2014) Protein interaction switches coordinate Raf-1 and MST2/Hippo signalling. Nat Cell Biol 16:673–84 doi:10.1038/ncb2986

57. Rose BA, Force T, Wang Y (2010) Mitogen-activated protein kinase signaling in the heart: angels versus demons in a heart-breaking tale. Physiol Rev 90:1507–46 doi:10.1152/physrev.00054.2009

58. Rössler U, Hennig AF, Stelzer N, Bose S, Kopp J, Søe K, Cyganek L, Zifarelli G, Ali S, von der Hagen M (2021) Efficient generation of osteoclasts from human induced pluripotent stem cells and functional investigations of lethal CLCN7-related osteopetrosis. J Bone Miner Res doi:10.1002/jbmr.4322

59. Sakai T, Naito AT, Kuramoto Y, Ito M, Okada K, Higo T, Nakagawa A, Shibamoto M, Yamaguchi T, Sumida T, Nomura S, Umezawa A, Miyagawa S, Sawa Y, Morita H, Lee JK, Shiojima I, Sakata Y, Komuro I (2018) Phenotypic Screening Using Patient-Derived Induced Pluripotent Stem Cells Identified Pyr3 as a Candidate Compound for the Treatment of Infantile Hypertrophic Cardiomyopathy. Int Heart J 59:1096–1105 doi:10.1536/ihj.17-730

60. Seabright M (1971) A rapid banding technique for human chromosomes. Lancet 2:971–972 doi:10.1016/S0140-6736(71)90287-X

61. Sheikh F, Raskin A, Chu PH, Lange S, Domenighetti AA, Zheng M, Liang X, Zhang T, Yajima T, Gu Y, Dalton ND, Mahata SK, Dorn GW, 2nd, Brown JH, Peterson KL, Omens JH, McCulloch AD, Chen J (2008) An FHL1-containing complex within the cardiomyocyte sarcomere mediates hypertrophic biomechanical stress responses in mice. J Clin Invest 118:3870–80 doi:10.1172/JCI34472

62. Shimizu M, Wang W, Walch ET, Dunne PW, Epstein HF (2000) Rac-1 and Raf-1 kinases, components of distinct signaling pathways, activate myotonic dystrophy protein kinase. FEBS letters 475:273–277 doi:10.1016/S0014-5793(00)01692-6

63. Strunk A, Bhalla V, Clopton P, Nowak RM, McCord J, Hollander JE, Duc P, Storrow AB, Abraham WT, Wu AH (2006) Impact of the history of congestive heart failure on the utility of B-type natriuretic peptide in the emergency diagnosis of heart failure: results from the Breathing Not Properly Multinational Study. Am J Med 119:69. e1–69. e11 doi:10.1016/j.amjmed.2005.04.029

64. Tohyama S, Hattori F, Sano M, Hishiki T, Nagahata Y, Matsuura T, Hashimoto H, Suzuki T, Yamashita H, Satoh Y, Egashira T, Seki T, Muraoka N, Yamakawa H, Ohgino Y, Tanaka T, Yoichi M, Yuasa S, Murata M, Suematsu M, Fukuda K (2013) Distinct metabolic flow enables large-scale purification of mouse and human pluripotent stem cell-derived cardiomyocytes. Cell Stem Cell 12:127–37 doi:10.1016/j.stem.2012.09.013

65. von Gise A, Lin Z, Schlegelmilch K, Honor LB, Pan GM, Buck JN, Ma Q, Ishiwata T, Zhou B, Camargo FD, Pu WT (2012) YAP1, the nuclear target of Hippo signaling, stimulates heart growth through cardiomyocyte proliferation but not hypertrophy. Proc Natl Acad Sci U S A 109:2394–9 doi:10.1073/pnas.1116136109

66. Wang L, Kim K, Parikh S, Cadar AG, Bersell KR, He H, Pinto JR, Kryshtal DO, Knollmann BC (2018) Hypertrophic cardiomyopathy-linked mutation in troponin T causes myofibrillar disarray and pro-arrhythmic action potential changes in human iPSC cardiomyocytes. J Mol Cell Cardiol 114:320–327 doi:10.1016/j.yjmcc.2017.12.002

67. Wang L, Kryshtal DO, Kim K, Parikh S, Cadar AG, Bersell KR, He H, Pinto JR, Knollmann BC (2017) Myofilament Calcium-Buffering Dependent Action Potential Triangulation in Human-Induced Pluripotent Stem Cell Model of Hypertrophic Cardiomyopathy. J Am Coll Cardiol 70:2600–2602 doi:10.1016/j.jacc.2017.09.033

68. Yang Y, Del Re DP, Nakano N, Sciarretta S, Zhai P, Park J, Sayed D, Shirakabe A, Matsushima S, Park Y, Tian B, Abdellatif M, Sadoshima J (2015) miR-206 Mediates YAP-Induced Cardiac Hypertrophy and Survival. Circ Res 117:891–904 doi:10.1161/circresaha.115.306624

69. Yu L, Daniels JP, Wu H, Wolf MJ (2015) Cardiac hypertrophy induced by active Raf depends on Yorkie-mediated transcription. Sci Signal 8:ra13 doi:10.1126/scisignal.2005719

70. Zhai Y, Luo Y, Wu P, Li D (2018) New insights into SERCA2a gene therapy in heart failure: pay attention to the negative effects of B-type natriuretic peptides. J Med Genet 55:287–296 doi:10.1136/jmedgenet-2017-105120

71. Zhao Z, Manser E (2015) Myotonic dystrophy kinase-related Cdc42-binding kinases (MRCK), the ROCK-like effectors of Cdc42 and Rac1. Small GTPases 6:81–88 doi:10.1080/21541248.2014.1000699

72. Zheng M, Dilly K, Dos Santos Cruz J, Li M, Gu Y, Ursitti JA, Chen J, Ross Jr J, Chien KR, Lederer JW (2004) Sarcoplasmic reticulum calcium defect in Ras-induced hypertrophic cardiomyopathy heart. Am J Physiol Heart Circ Physiol 286:H424–H433 doi:10.1152/ajpheart.00110.2003

73. Zhou Q, Li L, Zhao B, Guan KL (2015) The hippo pathway in heart development, regeneration, and diseases. Circ Res 116:1431–47 doi:10.1161/circresaha.116.303311

