## Supplemental data for "Alteration of myocardial structure and function in RAF1-associated Noonan syndrome: Insights from cardiac disease modeling based on patient-derived iPSCs"

**Abstract**
Noonan syndrome (NS), the most common among the RASopathies, is caused by germline variants in genes encoding components of the RAS-MAPK pathway. Distinct variants, including the recurrent Ser257Leu substitution in RAF1, are associated with severe hypertrophic cardiomyopathy (HCM). Here, we investigated the elusive mechanistic link between NS-associated RAF1S257L and HCM using three-dimensional cardiac bodies and bioartificial cardiac tissues generated from patient-derived induced pluripotent stem cells (iPSCs) harboring the pathogenic RAF1 c.770C>T missense change. We characterize the molecular, structural and functional consequences of aberrant RAF1 –associated signaling on the cardiac models. Ultrastructural assessment of the sarcomere revealed a shortening of the I-bands along the Z disc area in both iPSC-derived RAF1S257L cardiomyocytes, and myocardial tissue biopsies. The disease phenotype was partly reverted by using both MEK inhibition, and a gene-corrected isogenic RAF1L257S cell line. Collectively, our findings uncovered a direct link between a RASopathy gene variant and the abnormal sarcomere structure resulting in a cardiac dysfunction that remarkably recapitulates the human disease. These insights represent a basis to develop future targeted therapeutic approaches.

**Keywords:** Cardiac bodies, hypertrophic cardiomyopathy, iPSC, MAPK pathway, Noonan syndrome, RAF kinase, RASopathy, sarcomere

**Table S1.** The list of primers used in this study.

| Name | Sequence | Accession number |
|---|---|---|
| Actinin alpha 1 (ACTN1) | CCCGAGCTGATTGACTACGG | NM_001130005.1 |
| | GCAGTTCCAACGATGTCTTCG | |
| Alpha fetoprotein (AFP) | GAATGCTGCAAACTGACCACGCTGGAAC | NM_001354717.1 |
| | TGGCATTCAAGAGGGTTTTCAGTCTGGA | |
| Actin beta (ACTB) | AGCCTCGCCTTTGCCGA | NM_001101.5 |
| | CTGGTGCCTGGGGCG | |
| Calcium voltage-gated channel subunit alpha1 C (CACNA1C) | TGATTCCAACGCCACCAATTC | NM_199460.3 |
| | GAGGAGTCCATAGGCGATTACT | |
| Calcium voltage-gated channel subunit alpha1 D (CACNA1D) | CGCGAACGAGGCAAACTATG | NM_001128840.2 |
| | TTGGAGCTATTCGGCTGAGAA | |
| Troponin T2, cardiac type (TNNT2) | GGAGGAGTCCAAACCAAAGCC | NM_001001430 |
| | TCAAAGTCCACTCTCTCTCCATC | |
| Forkhead box A2 (FOXA2) | TGGGAGCGGTGAAGATGGAAGGGCAC | NM_021784.4 |
| | TCATGCCAGCGCCCACGTACGACGAC | |
| Hypoxanthine phosphoribosyl-transferase 1 (HPRT1) | CCTGGCGTCGTGATTAGTGAT | NM_000194.2 |
| | AGACGTTCAGTCCTGTCCATAAT | |
| Kruppel like factor 4 (KLF4) | CAGCTTCACCTATCCGATCCG | NM_001314052.1 |
| | GACTCCCTGCCATAGAGGAGG | |
| Myosin heavy chain 6 (MYH6) | GCCCTTTGACATTCGCACTG | NM_002471.3 |
| | GGTTTCAGCAATGACCTTGCC | |
| Myosin heavy chain 6 (MYH7) | ACTGCCGAGACCGAGTATG | NM_000257.3 |
| | GCGATCCTTGAGGTTGTAGAGC | |
| Myosin heavy chain 7B (MYH7B) | AGTGGCAATAAAAGGGGTAGC | NM_020884.4 |
| | CCAAGTTCACTCACATCCATCA | |
| Myosin light chain (MYL2) | TTGGGCGAGTGAACGTGAAAA | NM_000432.3 |
| | CCGAACGTAATCAGCCTTCAG | |
| Myosin light chain 7 (MYL7) | CATCAACTTCACCGTCTTCC | NM_021223.2 |
| | GAAGCTGCTTGAACTCATCC | |
| Nanog homeobox (NANOG) | CCCCAGCCTTTACTCTTCCTA | NM_024865.3 |
| | CCAGGTTGAATTGTTCCAGGTC | |
| Natriuretic peptide A (NPPA) | CAACGCAGACCTGATGGATTT | NM_006172.3 |
| | AGCCCCCGCTTCTTCATTC | |
| Natriuretic peptide B (NPPB) | TCCTGCTCCTGCTCTTCTTG | NM_002521.2 |
| | TCCTGTAACCCGGACGTTTC | |
| NK2 homeobox 5 (NKX2-5) | CCAGCCCTGCTCTCACG | NM_004387.4 |
| | GCCCAGCGTAGGCCTCT | |
| POU class 5 homeobox 1 (POU5F1)/OCT4 | CTGGGTTGATCCTCGGACCT | NM_002701.5 |
| | CACAGAACTCATACGGCGGG | |
| Phospholamban (PLN)/PLB | ACCTCACTCGCTCAGCTATAA | NM_002667.4 |
| | CATCACGATGATACAGATCAGCA | |
| Ryanodine receptor 2 (RYR2) | GGCAGCCCAAGGGTATCTC | NM_001035.2 |
| | ACACAGCGCCACCTTCATAAT | |
| ATPase sarcoplasmic/endoplasmic reticulum $Ca^{2+}$ transporting 2 (ATP2A2)/SERCA2 | CATCAAGCACACTGATCCCGT | NM_001681.3 |
| | CCACTCCCATAGCTTTCCCAG | |

| | | |
|---|---|---|
| SRY-box 2 (SOX2) | GCCGAGTGGAAACTTTTGTCG<br>GGCAGCGTGTACTTATCCTTCT | NM_003106.3 |
| SRY-box 17 (SOX17) | CGCTTTCATGGTGTGGGCTAAGGACG<br>TAGTTGGGGTGGTCCTGCATGTGCTG | NM_022454.4 |
| Titin (TTN) | CCCCATCGCCCATAAGACAC<br>CCACGTAGCCCTCTTGCTTC | NM_133378 |
| Troponin C1, slow skeletal and cardiac type (TNNC1) | TGGTTCGGTGCATGAAGGAC<br>GTCGATGTAGCCATCAGCATTT | NM_003280.2 |
| Troponin I3, cardiac type (TNNI3) | AGAAGGAGGACACCGAGAAG<br>GGAAGGCTCAGCTCTCAAAC | NM_000363.4 |
| Tubulin beta 3 class III (TUBB3) | ATGAGGGAGATCGTGCACAT<br>GCCCCTGAGCGGACACTGT | NM_006086.4 |
| Skeletal muscle (ACTA1)/α-SK (ACTA1) | TGCCAACAACGTCATGTCG<br>CAGCGCGGTGATCTCTTTCT | NM_001100.3 |
| Smooth muscle (ACTA2)/α-SMA | AAAAGACAGCTACGTGGGTGA<br>GCCATGTTCTATCGGGTACTTC | NM_001613 |
| Titin (N2B isoform) | GGCCGAGAAATTTATGAGAGTGAC<br>CGCTTTTCAGAACAACTTCTTCCT | NM_003319.4 |
| Titin (N2BA isoform) | CCAGCAACCAAGAAAGCTGCG<br>CCCAGAATCAGTTTTTGTGGTGTC | NM_001256850.1 |

**Table S2.** Reagents used in this study.

| REAGENT | SOURCE | IDENTIFIER |
|---|---|---|
| **Antibodies** | | |
| anti-OCT3/4 | Santa Cruz | #sc-5279 |
| anti-cardiac troponin t | Thermo Scientific | #MA5-12960 |
| anti-Tra-1-60 | Abcam | #ab16288 |
| anti-Myosin light chain 2 V | Synaptic Systems | #310111 |
| anti-SSEA4 | DSHB | #MC-813-70 |
| anti- γ -tubulin | Sigma-Aldrich | #T5326 |
| anti-phospho-ERK1/2 T202/T204 | Cell Signaling | #9106 |
| Anti-phospho-AKT S473 | Cell Signaling | #4060 |
| Anti-phospho-AKT T308 | Cell Signaling | #2965 |
| anti-phospho-YAP Ser127 | Cell Signaling | #4911 |
| anti-YAP | Cell Signaling | #4912 |
| anti-JNK | Cell Signaling | #9252 |
| anti-phospho-JNK Thr183/Tyr185 | Cell Signaling | #9251 |
| anti-S6K | Cell Signaling | #2708 |
| anti-phospho-S6K Thr389 | Cell Signaling | #9205 |
| anti-phospho-p38 Thr180/Tyr182 | Cell Signaling | #9211 |
| anti-p38 | Cell Signaling | #8690 |
| anti-alpha-actinin | Sigma-Aldrich | #A7811 |
| anti-ATP2A2/SERCA2 | Cell Signaling | #4388 |
| anti-RAF1 | Abcam | #AB181115 |
| anti-phospho-RAF1 S259 | Abcam | #ab173539 |
| anti-TUBB3 | Thermo Scientific | #MA1-118 |
| anti-Nkx2.5 | Santa Cruz | #sc-14033 |
| anti-Sox17 | R&D Systems | #AF1924 |
| anti-desmin | Agilent Technologies | #M076029-2 |
| anti-troponin I | Abcam | #Ab47003 |
| anti-SMA Clone 1A4 | DAKO | #M0851 |
| Custom made α-titin PEVK raised against PEVK S11878 (CEVVLKSVLRKR) and PEVK S12022 (LRPGSGGEKPP) (*Kötter, Sebastian, et al. Circulation research, 2016)* | Eurogentec | N/A |
| Anti-rabbit IgG Alexa Fluor 488 Conjugate | Cell Signaling | #4412 |
| Anti-mouse IgG Alexa Fluor 555 Conjugate | Cell Signaling | #4409 |
| Anti-mouse IgG Alexa Fluor 488 conjugated | Cell Signaling | #4408 |
| IRDye® 800CW Donkey anti-Rabbit IgG | LI-COR Biosciences | #926-32213 |
| IRDye® 800CW Donkey anti-Mouse IgG | LI-COR Biosciences | #926-32212 |
| Alexa488-conjugated goat anti-rabbit IgG | Thermo Scientific | #A11034 |
| Alexa546-conjugated goat anti-mouse IgG | Thermo Scientific | #A4671 |
| Alexa488-conjugated goat anti-mouse IgG | Thermo Scientific | #A11029 |
| **Chemicals, peptides, and recombinant proteins** | | |
| B-mercaptethanol | Sigma-Aldrich | #M-3148 |
| basic Fibroblast Growth Factor (bFGF) | Peprotech | #100-18B |
| ROCK-Inhibitor Y27632 | Selleckchem | #S1049 |
| CHIR99021 | Selleckchem | #S1263 |
| IWR-1 | Sigma-Aldrich | #I0161 |
| RPMI 1640 Medium, no glucose | Thermo Scientific | #11879-020 |
| human Albumin | Sigma-Aldrich | #A0237 |
| Sodium DL-Lactat | Sigma-Aldrich | #L4263 |
| L-Ascorbic acid-2-Phosphate | Sigma-Aldrich | #A8960 |
| DMEM / F12 + Glutamax | Thermo Scientific | #31331-028 |
| Knockout Serum Replacement (KO-SR) | Thermo Scientific | #10828028 |

| | | |
|---|---|---|
| MEM Non-Essential Amino Acids | Thermo Scientific | #11140-035 |
| RPMI 1640 Medium | Thermo Scientific | #21875-034 |
| B-27™ Supplement, minus insulin | Thermo Scientific | #A1895601 |
| B-27™ Supplement | Thermo Scientific | #17504044 |
| Geltrex Membrane Matrix | Thermo Scientific | #A1413201 |
| Gelatine from porcine skin | Sigma-Aldrich | #G2500 |
| Accutase™ Cell Dissociation Reagent | Thermo Scientific | #A1110501 |
| Collagenase, Type IV | Thermo Scientific | #17104019 |
| Agarose NEEO Ultra-Quality | CARL ROTH | #2267.4 |
| Hydrosil A and Hydrosil B | SILADENT | #101301 |
| AggreWell™400 | STEMCELL Technologies | #27840 |
| Versene Solution | Thermo Scientific | 15040066 |
| TrypLE™ Select Enzyme | Thermo Scientific | A1285901 |
| DMSO (dimethyl sulfoxide) | Sigma-Aldrich | #D2650 |
| Fetal Bovine Serum | Thermo Scientific | 26140079 |
| L-Glutamine | Thermo Scientific | 25030149 |
| KaryoMAX Colcemid | Thermo Scientific | 15212012 |
| Trypsin/EDTA | Biochrom | L 2143 |
| TRIzol™ | Thermo Scientific | 15596026 |
| SYBR™ Green | Applied Biosystems | #4309155 |
| Formaldehyde 4% | Carl Roth | #P087.1 |
| EDTA-free protease inhibitor | Sigma-Aldrich | 11873580001 |
| Phosphate buffered saline with 5% non-fat milk | Merck | #P4739 |
| Intercept® (TBS) Blocking Buffer | LI-COR | 927-60001 |
| ProLong™ Gold Antifade Mountant | Thermo Scientific | P10144 |
| Tissue-Tek® O.C.T. Compound | Sakura Finetek | #4583 |
| **Cell Lines** | | |
| HFF-1 | ATCC | SCRC-1041 |
| **Recombinant DNA** | | |
| pCE-hSK | Addgene | #41814 |
| pCE-hOct3/4 | Addgene | #41813 |
| pCE-hUL | Addgene | #41855 |
| pCE-mp53DD | Addgene | #41856 |
| pCXB-EBNA1 | Addgene | #41857 |
| **Oligonucleotides** | | |
| For primers, please see Table S1 | | |
| **Software** | | |
| FlowJo | Treestar, Ashland, OR | https://www.flowjo.com/ |
| IKAROS | MetaSystems (Altlußheim, Germany) | https://metasystems-international.com/de/products/ikaros |
| Image Studio 5.2 | LI-COR | https://www.licor.com/bio/image-studio-lite |
| ZEN 3.2 (blue edition) | Carl Zeiss AG | |
| IonWizard 6.4 | Ion Optix Corp | http://www.ionoptix.com/ |
| Prism 6 | GraphPad software | https://www.graphpad.com/scientific-software/prism/ |
| **Critical Commercial Assays** | | |
| DNA-free™ DNA Removal Kit | Invitrogen | AM1906 |
| GoScript™ cDNA Synthesis Kit | Promega | A5003 |

| | | |
|---|---|---|
| Quick Start™ Bradford Protein Assay | Bio-Rad | 5000201 |
| Trichrome II Blue staining kit | Roche | #860-013 |
| **Other** | | |
| 6-Well-Plate | Thermo Scientific | 140675 |
| T-25 Tissue Culture Flasks | TPP Techno Plastic | 90026 |
| T-175 Filter Cap Flasks | CELLSTAR | GR661175 |

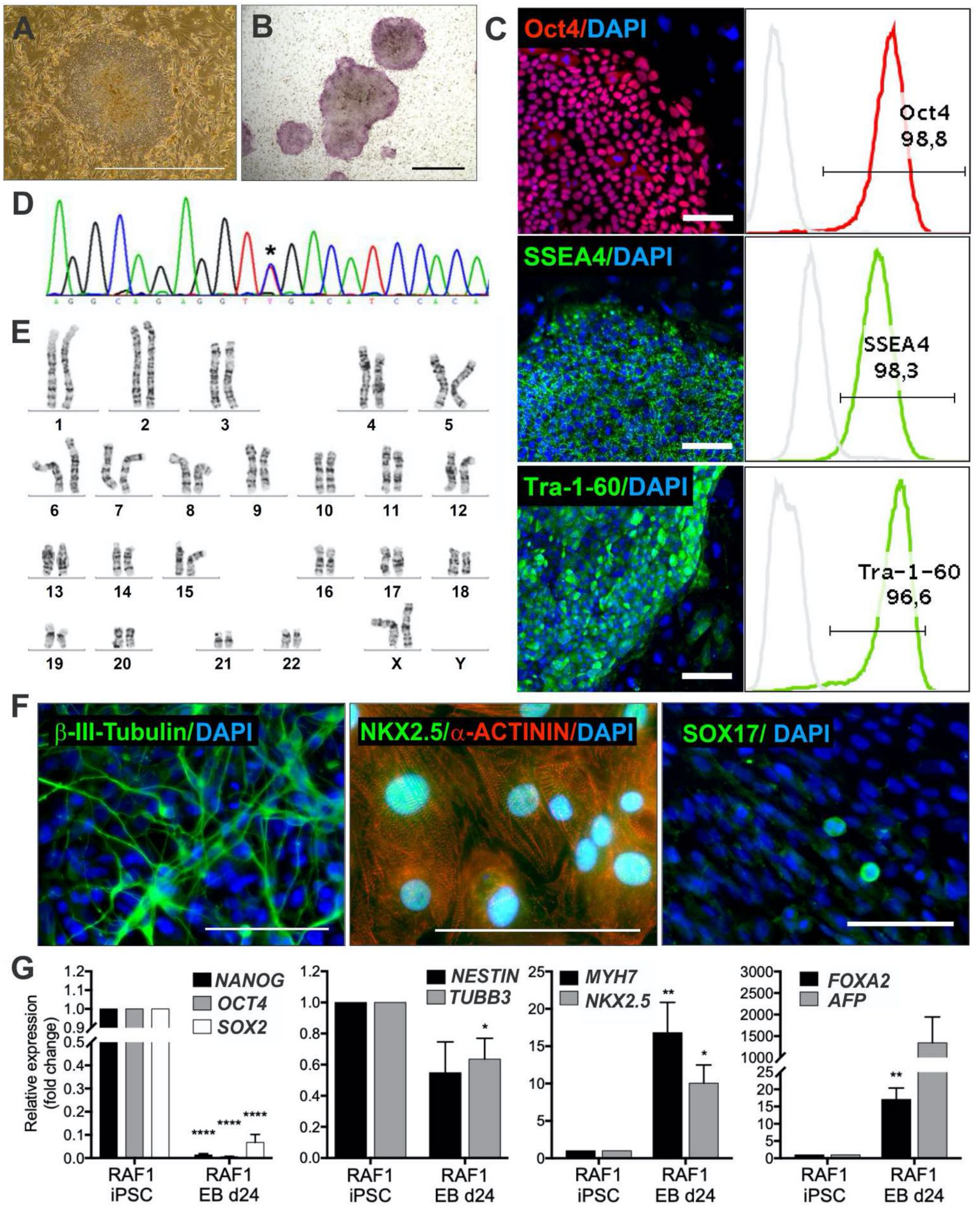

**Figure S1.** Characterization of RAF1$^{S257L}$ iPSC line 1 (7B10) from first patient.

A) Typical iPSC colonies on mitotically inactivated murine feeder cells. Scale bar, 1000 µm.

B) Alkaline phosphatase-positive colonies. Scale bar, 1000 µm.

C) Expression of pluripotency markers OCT4, SSEA4, and TRA-1-60 as detected by immunofluorescence staining and flow cytometry. Scale bar, 100 µm. Isotype controls of flow cytometry histograms depicted in light gray.

D) Sanger sequencing confirmed the heterozygous *RAF1 c.770C>T* variant in iPSCs (asterisk).

E) Normal diploid karyotypes in iPSCs at passage 8 after reprogramming.

F) Trilineage differentiation of patient derived iPSCs. Expression of ectodermal (beta-III-Tubulin), mesodermal (NKX2.5 and sarcomeric alpha-actinin), and endodermal (SOX17) markers was detected. Scale bars, 100 µm.

G) Relative gene expression of pluripotency (*NANOG*, *OCT4*, *SOX2*) and differentiation markers (*NESTIN*, *TUBB3*, *MYH7*, *NKX2.5*, *FOXA2*, *AFP*) of differentiated embryoid bodies on d24 of differentiation relative to undifferentiated iPSCs normalized by beta-ACTIN expression. Bar graphs represent mean of three independent samples +/- SEM. *P < 0.05, **P < 0.01, ***P < 0.001, ****P < 0.0001, unpaired t test. APF, alpha fetoprotein; EB, embryoid body; FOXOA2, forkhead box A2; MYH, myosin heavy chain; NKX2.5, NK2 homeobox 5; TUBB3, tubulin beta 3 class III, WT, wild-type.

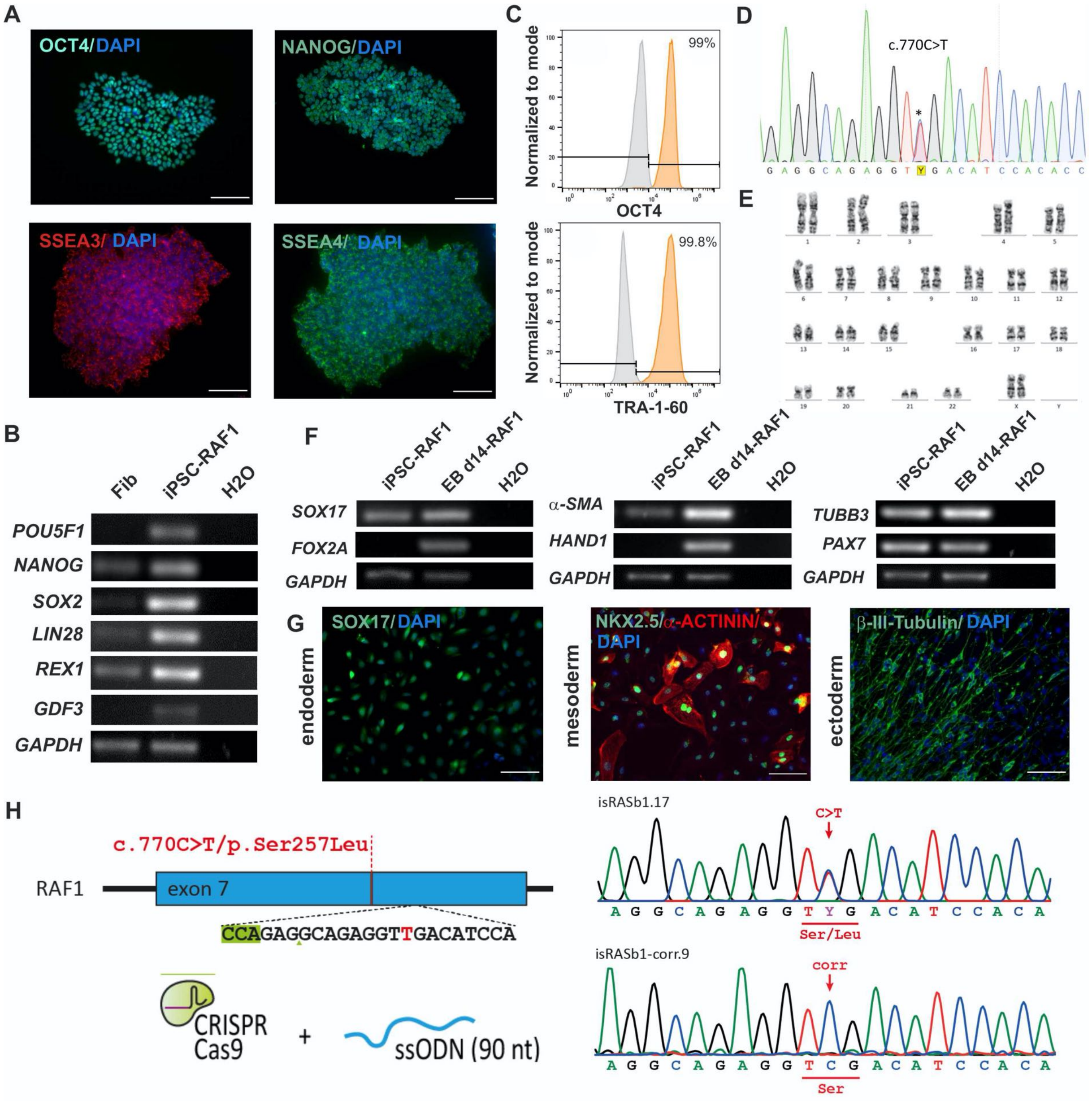

**Figure S2.** Characterization of RAF1$^{S257L}$ iPSC line 2 (isRASb1.17) from second patient. Human iPSC-RAF1$^{S257L}$ line 2 reveals expression of pluripotency markers, a normal karyotype, and differentiation potency towards ectodermal, endodermal and mesodermal derivatives *in vitro*.

A) iPSCs-RAF1$^{S257L}$ line 2 stain positive for OCT4, NANOG, SSEA3 and SSEA4 as detected by immunofluorescence staining. Scale bar, 200 μm.

B) RT-PCR analysis of pluripotency markers in iPSCs and control fibroblasts.

C) Flow cytometry analysis confirmed more than 99% of iPSCs stain positive for OCT4 and TRA-1-60.

D) Sanger sequencing confirmed the heterozygous RAF1$^{S257L}$ variant in iPSCs (asterisk).

E) iPSCs show a normal diploid karyotype.

F, G) Trilineage differentiation of iPSC-RAF1$^{S257L}$. Expression of endodermal (SOX17 and FOX2A), mesodermal (*α-SMA*, *HAND1*, NKX2.5, and sarcomeric alpha-actinin) and ectodermal (TUBB3 and *PAX7*), markers were detected by RT-PCR and immunofluorescence staining. Scale bar, 200 μm.

H) CRISPR-CAS9 mediated correction of the RAF1$^{S257L}$ mutation to RAF1$^{WT}$ for the line 2 iPSCs (isRASb1-corr.9).

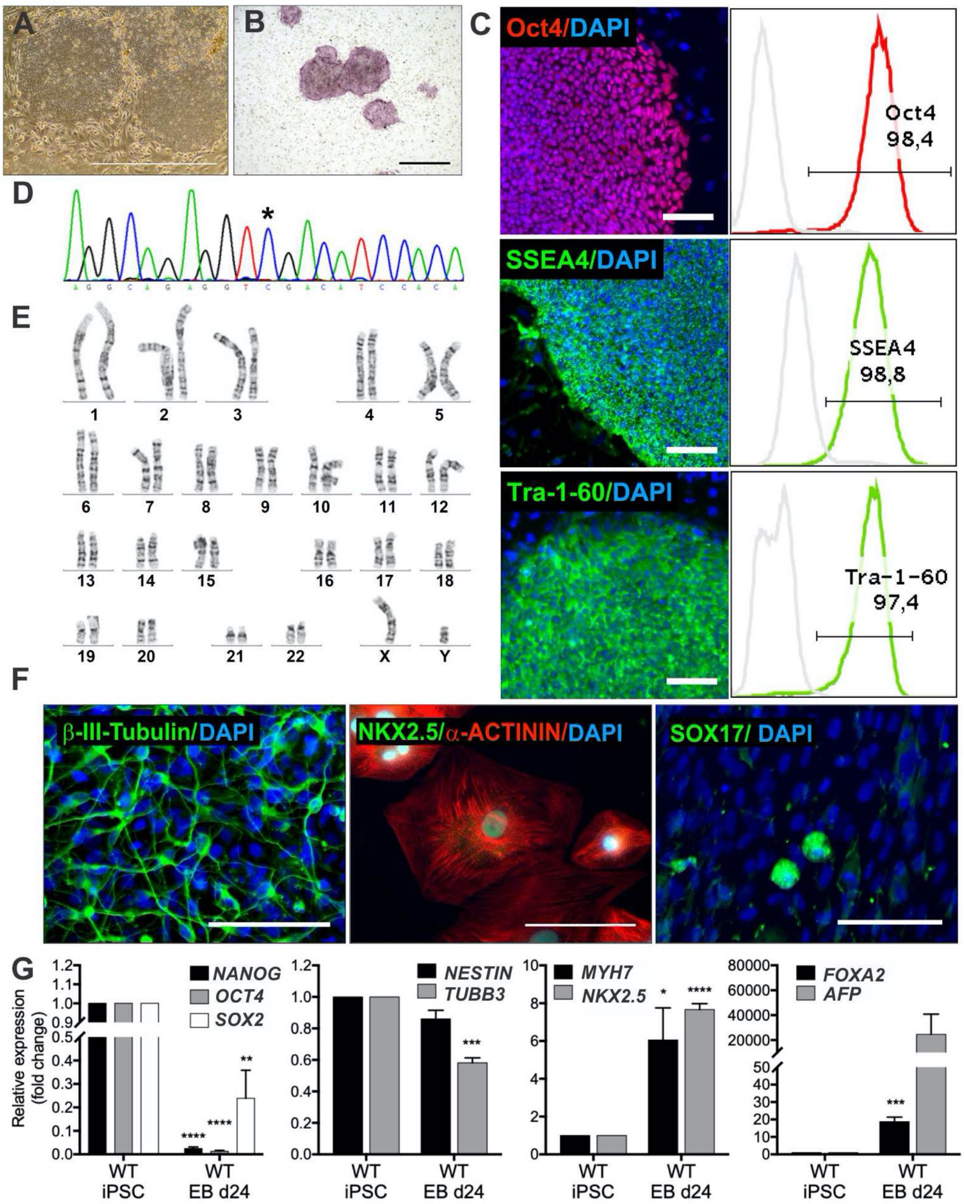

**Figure S3.** Characterization of WT iPSC line ipWT16.1 from donor 1 (WT1)
A) Typical iPSC colonies on mitotically inactivated murine feeder cells. Scale bar, 1000 µm.
B) Alkaline phosphatase-positive colonies. Scale bar, 1000 µm.
C) Expression of pluripotency markers OCT4, SSEA4, and TRA-1-60 as detected by immunofluorescence staining and flow cytometry. Scale bar, 100 µm. Isotype controls of flow cytometry histograms depicted in light gray.
D) Sanger sequencing confirmed the wildtype RAF1 sequence in iPSCs (asterisk).
E) Normal diploid karyotypes in iPSCs at passage 8 after reprogramming.
F) Trilineage differentiation of patient derived iPSCs. Expression of ectodermal (beta-III-Tubulin), mesodermal (NKX2.5 and sarcomeric alpha-actinin), and endodermal (SOX17) markers was detected. Scale bars, 100 µm.
G) Relative gene expression of pluripotency (*NANOG*, *OCT4*, *SOX2*) and differentiation markers (*NESTIN*, *TUBB3*, *MYH7*, *NKX2.5*, *FOXA2*, *AFP*) of differentiated embryoid bodies on d24 of differentiation relative to undifferentiated iPSCs normalized by beta-actin expression. Bar graphs represent mean of three independent samples +/- SEM. *P < 0.05, **P < 0.01, ***P < 0.001, ****P < 0.0001, unpaired t test. APF, alpha fetoprotein; EB, embryoid body; FOXOA2, forkhead box A2; MYH, myosin heavy chain; NKX2.5, NK2 homeobox 5; TUBB3, tubulin beta 3 class III, WT, wild-type.

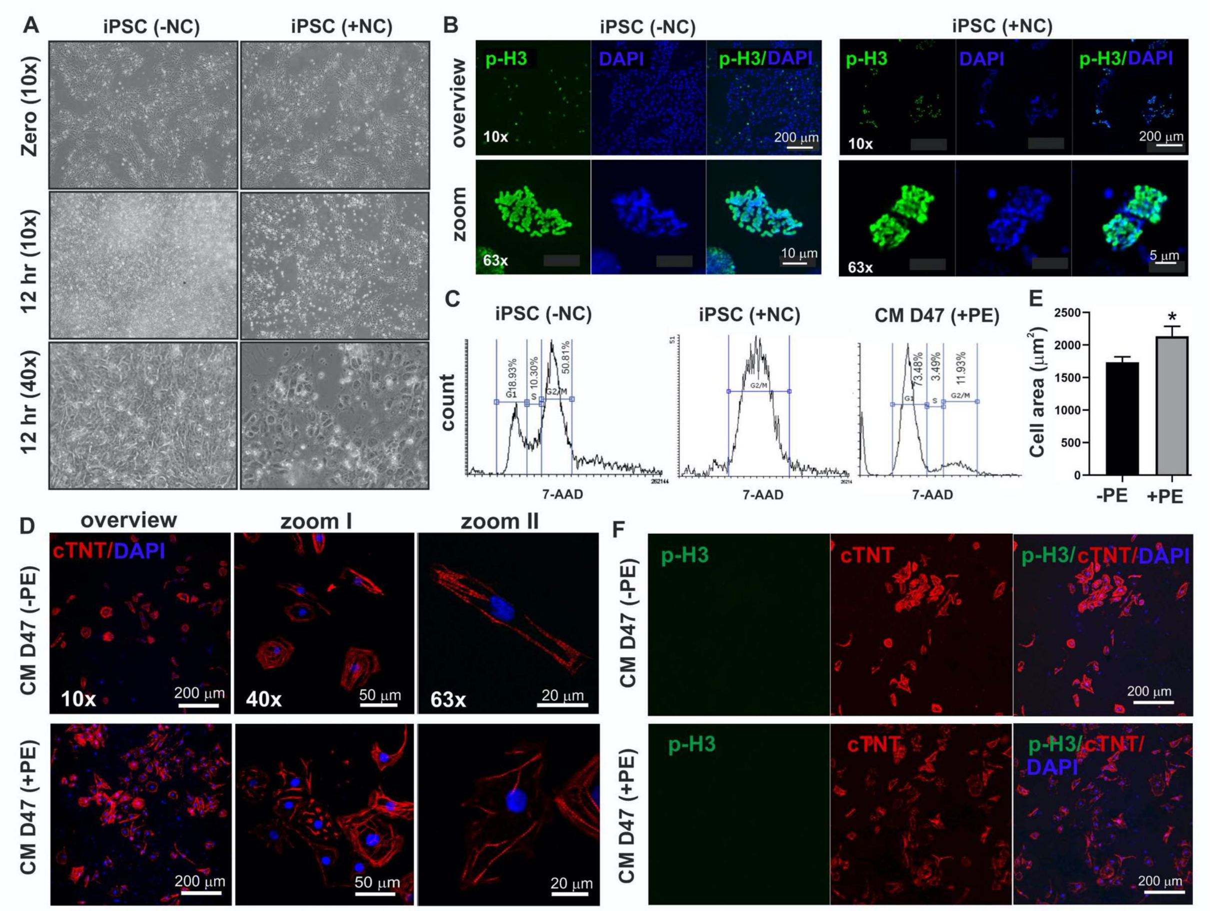

**Figure S4.** Morphology of the RAF1$^{S257L}$ iPSC line 1 after Nocodazole (NC) and phenylephrine (PE) treatments.

A) Illustration of cells arrested in mitosis upon NC treatment (100 nM) (lower panels).

B) phospho-histone 3 (p-H3) standing of human iPSC with different magnifications after NC treatment.

C) Cell cycle analysis of iPSC untreated and treated with NC as well as CMs treated with (PE).

D, E) cTNT staining of the dissociated CM showed increase in cell size of the CM treated with 100 nM of PE for 7-days. Confocal images of CMs treated with +PE were analyzed to determine the cell area using ImageJ software. Eighty-seven cells for the CTRL (-PE) and 59 cells for PE treatment were analyzed. The mean cell area is depicted and the SEM. Bar graphs represent mean of at least three independent samples +/- SEM. *P < 0.05, unpaired t test.

F) p-H3 staining of untreated (-PE) and PE-treated CM (+PE).

CM, cardiomyocytes; cTNT, cardiac troponin T; iPSC, induced pluripotent stem cells; PE, phenylephrine; p-H3, phospho-histone 3; NC, Nocodazole.

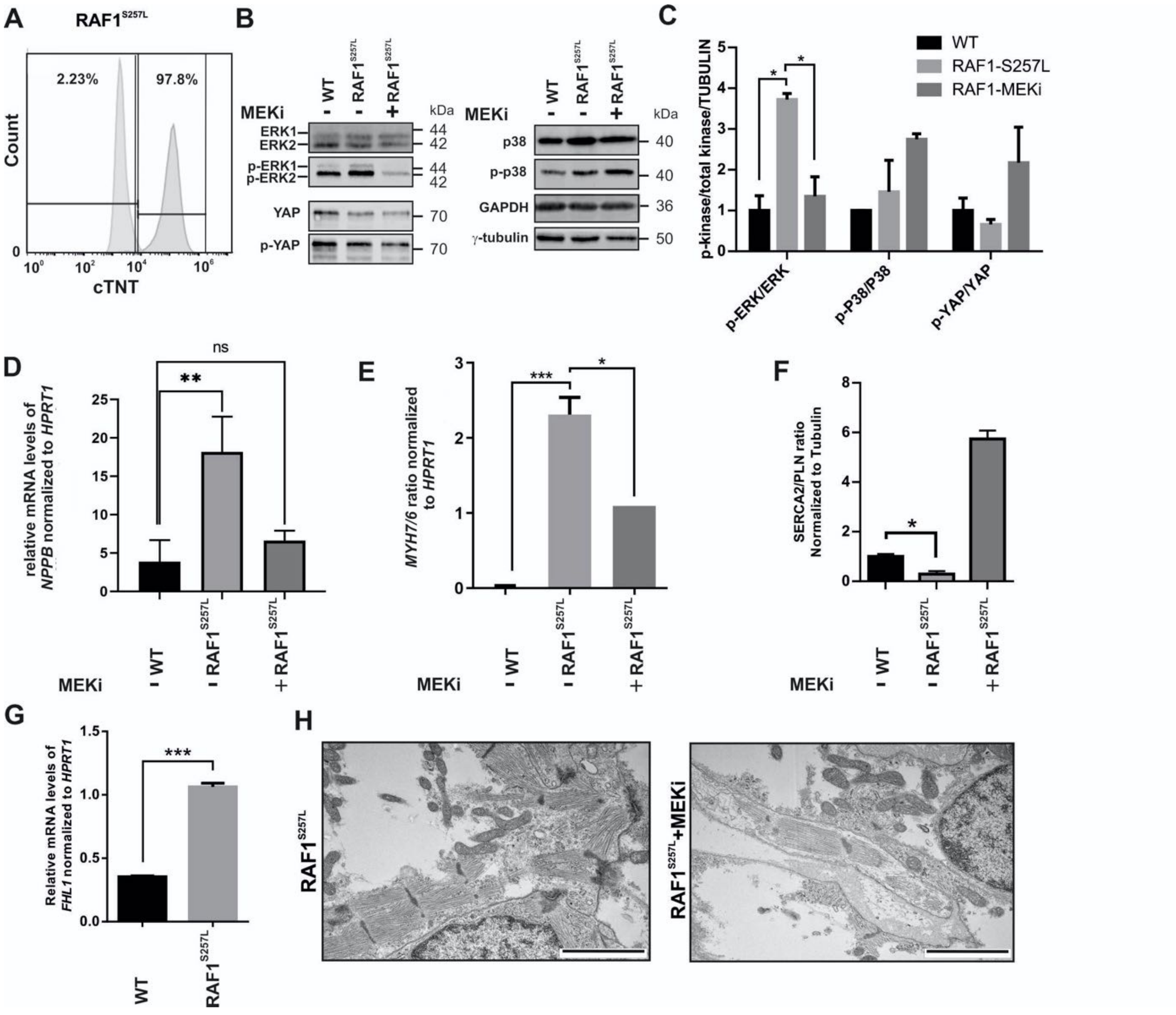

**Figure S5***.* Cardiac myocyte generation and analysis from second individual that carries RAF1$^{S257L}$.

A) Expression of cardiac troponin C marker by flow cytometry at day 24 of cardiac differentiation.

B) Western blots analysis of the mentioned signaling molecules, p-ERK/ERK, p-YAP/YAP, and p-p38/p38.

C) Quantification of phospho-protein to total protein ratios. *P < 0.05, unpaired 1-tail t-test, n≤2.

D) Quantitative real-time PCR analysis of *NPPB* transcript levels. **P < 0.01, unpaired 2-tail t-test, n≤2.

E) *MYH7-to-MYH6* ratio was compared between WT CBs and RAF1$^{S257L}$ CBs according to $2^{(-\Delta ct)}$ values.

F) Quantification of western blot results of SERCA2/PLN protein ratio normalized to Tubulin. *P < 0.05, unpaired 2-tail t-test, n≤2.

G) Quantitative real-time PCR analysis of *FHL1* transcript levels, n≤2, technical replicates.

H) Electron microscopic images of RAF1 mutated CBs treated with 0.2 μM MEK inhibitor from d12 of differentiation, scale bar, 2 μm.

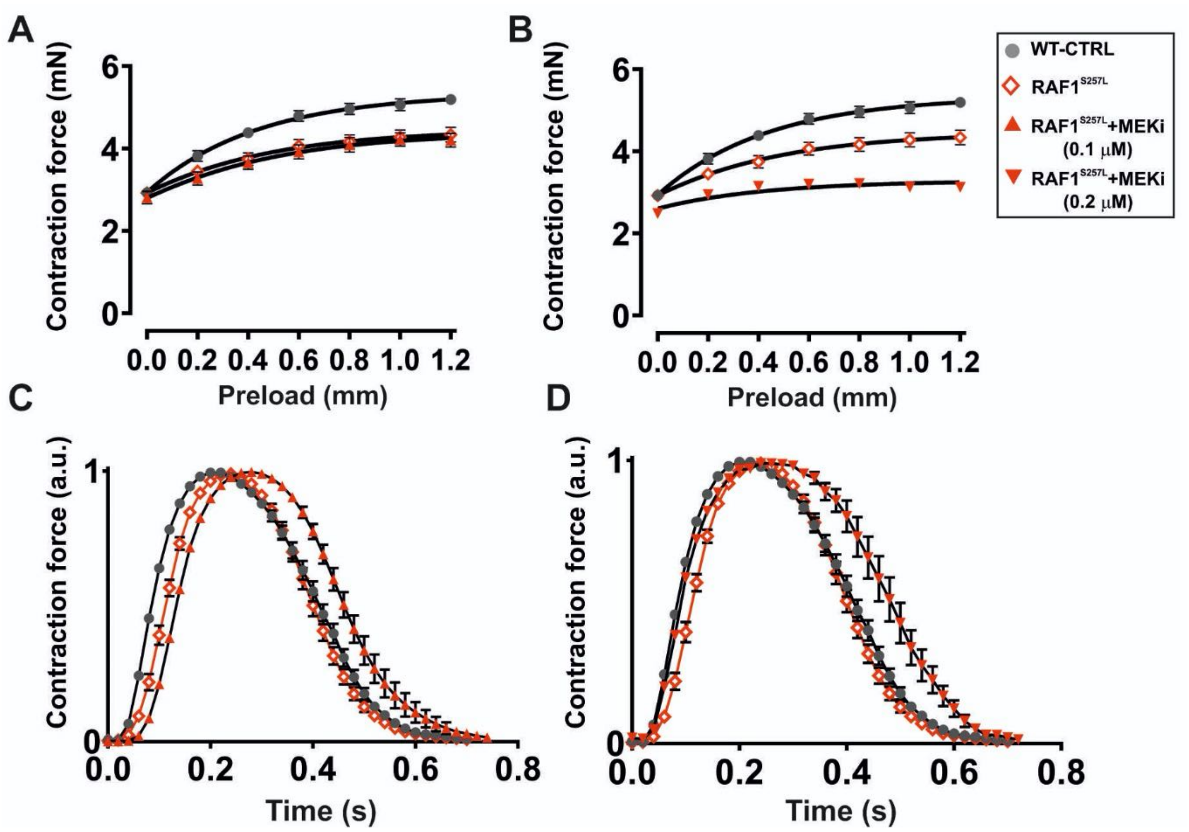

**Figure S6.** Effect of MEKi on the force development of human iPSC-derived bioartificial cardiac tissue.
A) Frank-Starling mechanism of: WT + RAF1 w/o + RAF w/ 0.1 µM MEKi.
B) Frank-Starling mechanism of: WT + RAF1 w/o + RAF w/ 0.2 µM MEKi.
C) Normalized contraction of: WT + RAF1 w/o + RAF w/ 0.1 µM MEKi.
D) Normalized contraction of: WT + RAF1 w/o + RAF w/ 0.2 µM MEKi.
